# Developing *SCL2205*: A Protein Sequence-based Spatial Modelling Dataset for the Protein Language Model Frontier

**DOI:** 10.64898/2026.03.08.710388

**Authors:** Daniel Ouso, Gianluca Pollastri

**Affiliations:** School of Computer Science, University College Dublin, Dublin 4, Belfield, Dublin, Ireland; Centre for Research Training in Genomics Data Science, University of Galway, University Rd, Galway, H91 TK33, Ireland

**Keywords:** Subcellular localisation, Machine learning, Protein Language Models, Data leakage, Data augmentation, Sequence modelling

## Abstract

Deep learning (DL) has advanced computational genome annotation tasks such as protein sub-cellular localisation (SCL) prediction. Nonetheless, its potential remains underutilised, primarily because of the limited availability of high-quality reference data and suboptimal input preparation strategies. In this study, we develop and analyse a high-quality dataset derived from the latest release of the universal protein knowledgebase (UniProtKB), designed to address existing challenges and support robust DL-based SCL modelling. The dataset was constructed through extensive quality preprocessing to ensure reliability, manual label mapping to enhance the quantity and diversity of the training data, and stringent partitioning to minimise data leakage. We validated the dataset using independent test sets, achieving up to 10.8% performance improvement, measured by the area under the precision–recall curve (PR-AUC), compared to the state-of-the-art (SoTA). Furthermore, we highlighted potential performance metric inflation in existing SoTA predictors by demonstrating, for the first time, at least 4.8% *training-to-testing* data leakage (pre-sequence representation) when using only 10% of the training set under homology augmentation (augmentation based on sequence similarity database searches; details in **Sub-section 2.1**), a commonly used data augmentation strategy in DL-based SCL prediction modelling. *SCL2205* will efficiently support the development of robust, trustworthy, and generalisable DL-based SCL predictors, while minimising data leakage and promoting reproducibility. It is openly available under the Creative Commons Zero (CC0 1.0) licence on DRYAD and is conveniently deployed as a package on the Python Package Index – `p-scldata`.

## Introduction

Automated sequence annotation is a critical component of functional genomics. It is mainly driven by widespread access to massively parallel sequencing. The adoption of Artificial intelligence (AI) for protein cellular component location annotation remains an active research area currently dominated by sequence-based DL predictors. Owing to its ability to model complex phenomena and directly utilise sequence information, DL has become more popular than classical machine learning (ML). The latter approach relies on pre-determined sequence characteristics as modelling features. However, sufficient quality training data is the main bottleneck to efficiently leveraging supervised DL predictors, which are very data-intensive. Regardless of the data challenge, it is paramount to provide high-quality training input.

Although UniProtKB is the primary source of sequences for SCL DL predictors, researchers often prepare their data differently, introducing avoidable bias. These biases may compromise the fair and accurate evaluation of different predictors. Moreover, suboptimal handling of intrinsic yet problematic data characteristics, such as sequence homology, can lead to costly consequences, such as training-to-testing data overlap. In addition, some predictors continue to rely on outdated database versions, despite significant improvements in data quality and quantity. This is likely associated with the rigorous and time-consuming nature of data development.

Notable advancements in AI have been driven by well-curated datasets across domains, including computer vision [19], natural language processing [20], and protein structure prediction – for example, the Critical Assessment of Structure Prediction (CASP) datasets. In the SCL domain, the DeepLoc dataset [3] is well known, alongside others such as MultiLoc [15]. In DeepLoc, sequences were filtered to include only eukaryotic, non-fragment, nucleus-encoded proteins, with a minimum length of 41 amino acids and confirmed experimental annotations [3, 29]. In contrast, MultiLoc [15, 5] applies filters based on eukaryotic origin and selected keywords from sequence comments and features. Given the processing disparities, which make performance comparisons problematic, it would be desirable to standardise and consistently curate the input data across all predictors.

As concerns emerge on sustainability and the environmental costs of training DL-based AI, the need for high-quality training input is critical to mitigate data noise that inflates training costs. Consequently, recent research has demonstrated gains with small yet well-curated training datasets in leading domains of AI such as large language models (LLMs) [21]. On the other hand, trustworthiness and the precision of the performance report in AI research are common concerns, especially in critical sectors such as biology and health. Similarly, compliance efforts are proposed for biological AI modelling [31]. In light of the above, we set out to develop a robust SCL dataset for DL predictor modelling, whilst underscoring current challenges and positioning for emerging frontiers. Beyond common preprocessing practices like filtering, we make the following major contributions with our dataset:

1. We highlight the limitations of homology augmentation by affirming and, for the first time, quantifying its contribution to data leakage using metrics comparable to those used in homology reduction, thereby underscoring trustworthiness in predictors.
2. Unlike in other cases, we perform extensive and robust homology reduction by minimising training-to-validation and training-to-testing sets overlap to *≤* 30% homology, while preserving the sequence-length distribution of the original dataset.
3. We provide two streams of the dataset – *SCL2205* – for model training: (i) a *training-validation* split and (ii) a *five-fold cross-validation* set. In addition, a *held-out* independent testing set is provided for final evaluation.
4. We employ domain knowledge to manually map labels to higher-level cellular components commonly used with SCL predictors.

We propose a novel comprehensive dataset (*SCL2205*) for application in SCL modelling, unlike the minimally described ones in most SCL models. For reproducibility, we discuss in detail the processes and design choices used to generate the dataset. This work establishes a new benchmark dataset that will empower researchers to build more sustainable, precise, and trustworthy tools for the next frontier of AI in spatial genomic discovery.

## Methodology

### Terminology

#### Sequence homology

According to Medical Subject Headings (MeSH), it is the degree of similarity between sequences. The similarity is attributed to descent from a common ancestor and can be based on percentage sequence identity and/or percentage positive substitutions. While similarity within diverse sets is important in modelling, class over-representation can be problematic. Mitigation is needed to avoid over-weighting closely related sequences. In database parlance, sequence homology is commonly synonymous with *redundancy*; therefore, redundancy reduction is associated with homology mitigation.

#### Homology reduction

It is the process of minimising sequence homology across or within datasets. In this study, we distinguish *across* and *within* dataset homology reduction by as *overlap* and *redundancy*, respectively.

#### Data leakage

Refers to data overlap across independent datasets. When in excess, it is undesirable in model training, where sufficient separation is needed across the *training-validation* and *training-testing* sets. The separation serves two major purposes: (i) enabling accurate model performance assessment and (ii) ensuring model inferential robustness to unseen data.

#### Data augmentation

Data augmentation increases the size and/or enriches the quality of available data, aiming to enhance model performance. It helps mitigate the problem of minimal training data, especially for supervised learning. The details of data augmentation are elaborately covered elsewhere [26, 27, 10]. From a broad perspective, it may be categorised into three approaches: (i) input-level augmentation, which involves transformations applied in the data space; (ii) embedding-level augmentation, which operates in the feature space and targets intrinsic relationships and semantics; and (iii) hybrid augmentation, which combines elements of both input- and embedding-level strategies. Alternatively, from the perspective of the augmentation source, methods may be classified as either *internal* or *external*. Internal augmentation refers to modifying the existing data directly, for example, reordering elements within a sequence. In contrast, external augmentation involves incorporating additional information, such as generating sequence profiles through sequence alignments. Although protein SCL prediction borrows much from the text domain of sequential learning, it has unique requirements and/or characteristics. Therefore, most of the natural language processing (NLP) augmentation techniques are not (yet) implemented in the SCL modelling domain. However, since the latter is affected by imbalanced and limited labelled data, other augmentation techniques are applied, mainly, homology augmentation, yet not without inherent limitations, especially since it counteracts homology reduction. Homology augmentation can be categorised as embedding augmentation, while label mapping falls under input augmentation.

#### Homology augmentation

Homology augmentation refers to sequence data enhancements involving database searches to retrieve additional sequences related to the original dataset. The additional sequences can be directly included in the original set or used to build sequence profiles from multiple sequence alignment (MSA). The latter is common with DL SCL predictors.

### Data Source

#### Ethical considerations

No ethical pre-approval was required. The raw data were retrieved from an open, public protein sequence database– UniProtKB, which does not contain any personal information.

#### The Universal Protein Knowledgebase

The UniProtKB is the central repository for aggregating functional information on proteins characterised by consistent, accurate and rich annotations. The annotations include universally accepted cross-references, ontologies, classifications, and quality scores informed by experimental and computational evidence [7]. The knowledge-base consists of two sections: (i) UniProtKB/Swiss-Prot for manually-annotated records based on extracted information from literature and expert-evaluated computational analysis, and (ii) UniProtKB/TrEMBL for computationally analysed records awaiting full manual annotation [7]. We used the former source because of the quality assurance of its records. More than 95% of proteins in UniProtKB are derived from translations of coding sequences (CDS) submitted to the International Nucleotide Sequence Database Collaboration (INSDC) databases [7]. In UniProtKB/Swiss-Prot, all protein products encoded by a gene in a given species are represented in a single record; that is, identical sequences are captured as a single record. However, if the sequences are from different species, they form different records [7].

#### Protein subcellular location labels

Most SCL predictors are trained with *supervision*. The *Subcellular location* subsection of UniProtKB contains the protein cellular location annotations used to supervise training. The UniProtKB provides location and topology information for the mature protein in the cell using a structured hierarchy of controlled vocabulary, except for the *Note*, which is free text [7]. Because proteins may be of mitochondrial, plastid or nucleic origin, it is important to note that the label annotations represent the mature protein, that is fully processed and functional, resulting from post-translation modification (PTM) and proteolytic processing. It is the ultimate version associated with a biological function. Therefore, filtering sequences on genomic origin (mitochondria/plastid/nuclear) [3, 29] may be unnecessary in general predictor training.

#### Raw data

Protein sequences and their annotation information were manually retrieved in tabular format from the latest release of the *Reviewed* subset of UniProtKB (UniProtKB/Swiss-Prot, Release 2022_05; date: 2023-01-24). Each record includes a stable and unique *Entry* identifier, which is essential for provenance and reproducibility.

### Data inclusion criteria

A total of 469,935 sequence records were downloaded. The following five filtering processes were applied; *data field name: process description*:

1. *Sub-cellular location [CC]*: Records lacking SCL annotation removed.
2. *Sub-cellular location [CC]*: Annotations with an evidence code ontology (ECO) code of *ECO:0000269* for the evidence – annotation is experimentally determined – were retained. These annotations are a credible basis for supervisory learning.
3. *Taxonomic lineage*: Retained records belonging to the eukaryotic taxonomic group using *Eukaryota (super-kingdom)* as the match phrase in the description. The eukaryotic cell is very similar across species; therefore, we minimise data fragmentation across taxonomic *Kingdoms* by aggregating at the super-kingdom level.
4. *Annotation*: Annotations with a quality score of at least three were retained. Although often overlooked, the cut-off enforces stringent reference data quality to minimise compounding errors.
5. *Length*: Sequences with at least 30 and at most 5,000 amino acids. Evidence shows biological function in shorter sequences; however, most SCL predictors cap sequence length from 30 to 40 amino acids.

Most current SCL datasets stop at some of the above minimal preprocessing before proceeding to the homology reduction step. However, we harness more from the data than in earlier research. The DeepLoc dataset [3, 29] incorporates label mapping for various locations. However, it is minimal, based on an older database release and may contain some avoidable noise. For instance, while organellar-membrane-localising proteins are mapped to their respective organelles, membranes, relative to the organelles, are structurally and functionally related across organelles, in a general sense.

Therefore, it might be better to consider them as a location entity, at least for a general SCL predictor, rather than organelle-specific. The latter approach is adopted in some membrane protein predictors: when predicting endomembrane system and secretory pathway proteins [18] or membrane and non-membrane proteins [17].

#### Label mapping

In the next section, we exploit the biological controlled vocabulary SCL definitions to map sequences of rare sub-compartment SCL labels, which would otherwise be discarded, to their higher-order compartments; the labels/locations we are interested in predicting. The intuition is simple: for a sequence to belong in a *sub-space*, it must belong in a *space*. Because *Membrane* is a unique entity, sub-membranes (membranes within a compartment) were lump-summed under *Membrane*. Therefore, the granularity, or the predicted location of interest, matters more and is partly determined by the amount of representation for that location.

Mapping began by first extracting and restructuring label information from the structured annotation text using custom code. The resulting labels are concise, sorted lists containing only the unique SCL annotation(s) for a sequence. Afterwards, manual label mapping using UniProtKB’s SCL ontology information [7] and the interactive cell map from *SwissBioPics* library [22] as reference ensued. We mainly considered rare labels with five or more sequences for mapping.

For completeness, *single-location* and *multi-location* proteins were included in the final dataset (*SCL2205*). However, for simplicity, only the *Single-location* proteins were used for experiments. **Fig 1** illustrates the process of label mapping. Ultimately, the process delivered a substantial increase in sequence count (see **Table 1**), which is valuable for training.

**Table 1.** A summary of the sub-cellular location (SL) composition showing the mapping effect. The first and second columns are the *before* and *after* protein counts, respectively. The fold increase is captured in the last column. Overall, the sample size grew by 71%. When considering only the single-location proteins, the numbers improved by 80%. For brevity, *Centrosome; Cytoplasm; Cytoskeleton; Microtubule organising center* is shortened as *Cent; Cytop; Cytos; MTOC*.

| Label | Original | Mapped | Fold Increase |
| --- | --- | --- | --- |
| Membrane | 290 | 5616 | 19.37 |
| Nucleus | 3621 | 4721 | 1.30 |
| Secreted | 3067 | 3357 | 1.09 |
| Cytoplasm | 2209 | 2706 | 1.22 |
| Cytoplasm; Nucleus* | 2167 | 2618 | 1.21 |
| Mitochondrion | 838 | 1062 | 1.27 |
| Plastid | 6 | 622 | 103.67 |
| Cent; Cytop; Cytos; MTOC* | 92 | 338 | 3.67 |
| Cytoplasm; Membrane* | 88 | 284 | 3.23 |
| Cytoplasm; Cytoskeleton* | 244 | 270 | 1.11 |
| ER | 150 | 204 | 1.36 |
| Cell projection | 6 | 194 | 32.33 |
| Peroxisome | 144 | 160 | 1.11 |
| **Totals** | **12922** | **22152** | **1.71** |

\* Multiple-localisation.

**Fig. 1.**
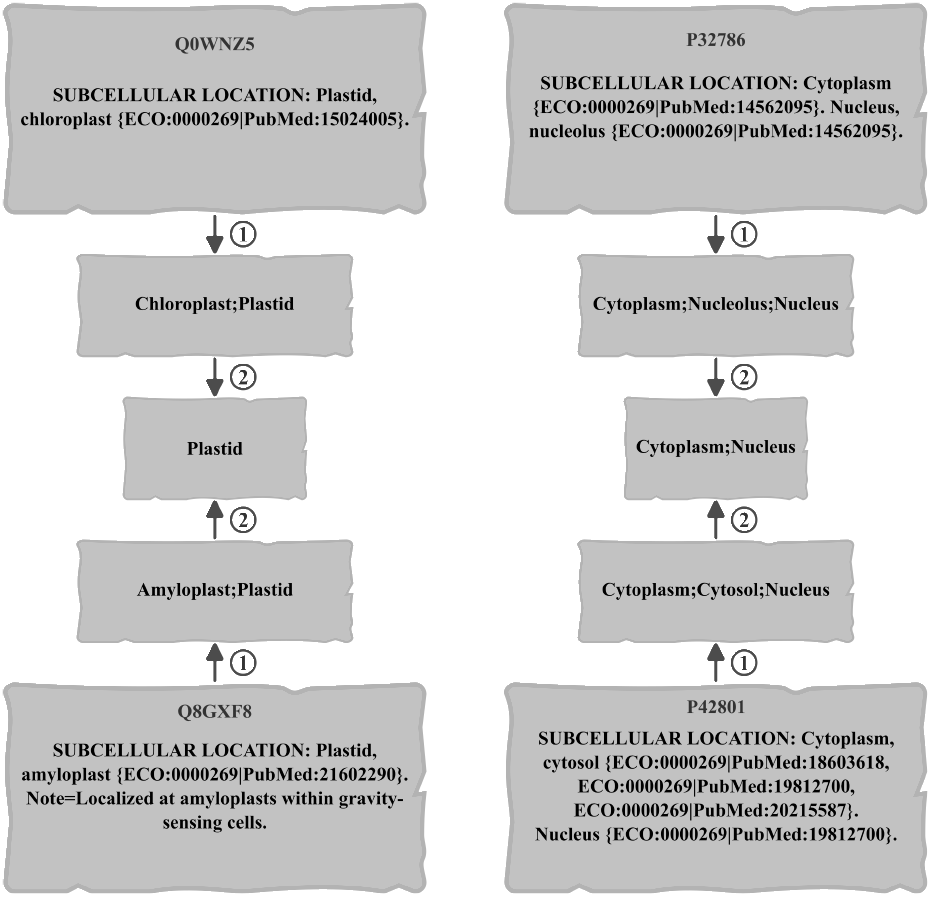
Illustrating SCL label mapping using four proteins. The left and right panels represent *single-location* and *multi-location* protein mapping, respectively. **Step 1**: programmatically, extract labels from structured annotation text using custom code. **Step 2**: manually maps sub-location (sub-compartment) to location labels and harmonises interchangeable labels. **Alt text**: Graphics illustrating the mapping processes using a pair of single-localisation and multi-localisation proteins; involving mining labels and manual mapping.

#### Homology reduction

Essentially, homology reduction is common in protein modelling for two reasons: to mitigate data leakage and avoid learning bias; it mainly relies on sequence alignment. The *CD-HIT* tool [11], which is based on position-specific iterative BLAST (PSI-BLAST), is widely used by DL predictors to facilitate homology reduction. *CD-HIT* prefers longer sequences, despite short and long sequences having functional roles. However, at least two issues arise: unnatural input data distribution and loss of functional domain (pattern) representation that could be useful for learning SCL. Consequently, we implemented a custom variant sequence similarity algorithm, which is still based on basic local alignment sequence tool (BLAST), factors sequence-pair length, but without preference for longer sequences, and conveniently operates on *training-testing* split dataset models for ML; avoiding multi-stage homology reduction regardless of the homology threshold. The similarity algorithm is detailed in the **Methods** section of **Supplementary File S1**.

Based on the reasons already provided, we had the following goals for our homology reduction pipeline (**Fig 2**):

**Fig. 2.**
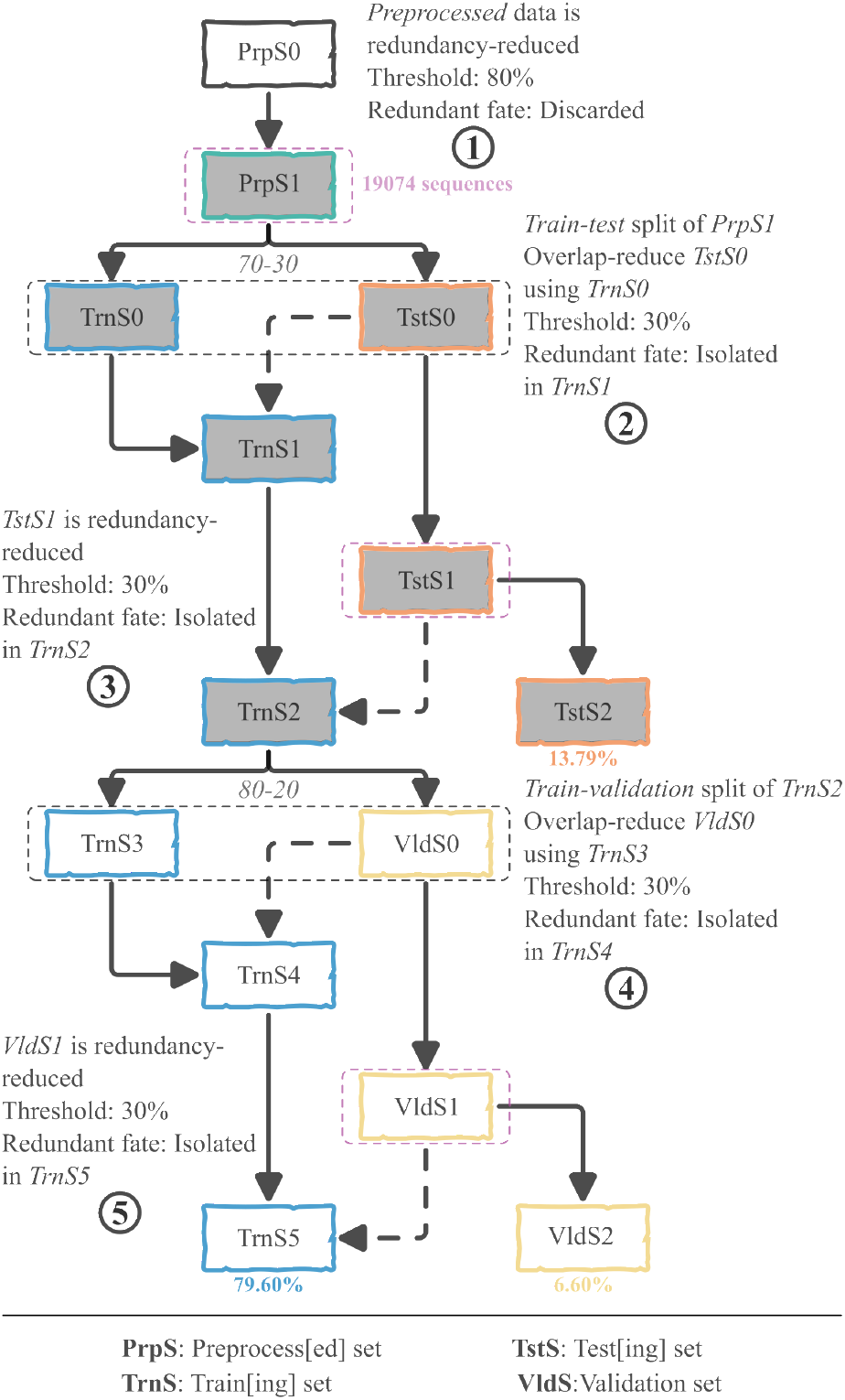
Summary illustration of the three-step homology reduction process based on global pairwise alignments. Solid-line arrows indicate data processing flows, broken-line arrows represent data isolation, and bifurcating arrows are data splits (split ratios are included at the nodes). Magenta, dashed rectangles denote redundancy reduction, while the black ones correspond to overlap reduction. The shaded rectangles highlight the repeating *motif* within the homology reduction pipeline. The process takes the 22,152 preprocessed (*PrpS0*) instances as input. It then begins from a global redundancy reduction that results in 19,074 instances ((*PrpS1*; 100%), resulting in 15,183 instances in the training set (*TrnS5*; 79.60%), 1,260 in the validation (*aka* discovery or development) set (*VldS2*; 6.60%), and 2,631 as the held-out testing set (*TstS2*; 13.79%). **Alt text**: Graphic illustrating the three-step homology reduction process. Redundancy reduction occurs within datasets, while overlap reduction occurs across dataset partitions.

1. **Mitigating data leakage** – similar samples stay in a bin; *training* or *testing*, not both.
2. **Avoiding learning bias; imbalance, by implication** – maintain a particular similarity threshold within a bin.

We then proceeded with a three-step homology reduction strategy based on similarity thresholds and dataset partitions. In the first step, we minimise overweighting associated with imbalanced classes and high sequence homology during training by using an 80% similarity threshold to redundancy-reduce the preprocessed dataset (**Fig 2**, *PrpS0*). The second and third steps use 30% similarity threshold while considering the partitioned data. The testing set (*TstS0*; subject; 30%) was overlap-reduced against the training set (*TrnS0*; query; 70%) and the testing set (subject and query) redundancy-reduced, respectively. Whereas the second step mitigates data leakage, the third step mitigates evaluation bias, which is associated with imbalanced classes and sample similarity. The overlapping and redundant testing set sequences were isolated (binned; **Fig 2** broken arrows) in the training set to align with the goals.

Moreover, we used the resulting training set to generate two dataset tracks for model training, based on a common model development data partitioning approach: *training, validation*, and *testing*; and *cross-validation* and *testing*. Henceforth, we refer to the two tracks as *training-validation-testing* (TVT) and *cross-validation-testing* (CVT), respectively, and collectively call them *SCL2205*. Note that the held-out testing set is shared because the original dataset is the same. Therefore, onwards we refer to the partitions associated with the model training process as *training-validation* (TV) and *cross-validation* (CV). As before, the TV and CV sets were subjected to the second and third steps of homology reduction. This was across five folds for the CV set. In the TV set, the *training* (**Fig 2, *TrnS3***) and *validation* (**Fig 2, *ValS0***) sets were split at 80% and 20% respectively.

##### Note

To ensure reproducibility, all data partitioning was performed using the `train_test_split` function from the *scikit-learn* library, with a consistent random seed of 2023 maintained throughout the study. Other software details are in **Supplementary File S1**.

## Experiments

### Manual mapping impact

We aimed to assess the contribution of manual label mapping on model generalisation, compared to using the native labels. Using *SCL2205*_*s*_ as the reference, we generated 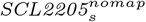 by copying *newData*_*s*_ and removing all sequences associated with the label-mapping augmentation process. Since very few samples in the *Cell projection* and *Plastid* categories remained when considering the original labelling, we omitted them from the modelling process. The corresponding partitions of the sets were used to train two convolutional neural network (CNN)-based networks as later outlined in **Sub-subsection 3.3.2**. The independent test sets were redundancy-reduced per the two datasets and used for evaluation.

### Implication of data leakage from homologous augmentation

How does homology augmentation contribute to and impact the overlap between *training-testing* datasets, especially for predictors based on pre-model-training homology reduction?

To attempt answering the question, we designed an experiment using the TVT dataset. Whilst maintaining class proportions, we randomly selected 10% (*n* = 1280; *random-training-seqs-A*) sequences of the training set (**Fig 2**; *TrnS5* minus multi-localising sequences; 12,807) sequences and queried the *RefSeq* (Version 5 of 202410260537) database for homologues using BLAST global alignment – as is often the case. The tweaked PSI-BLAST parameters were: `evalue=0.001; num_iterations=3; matrix=BLOSUM62, word_size=3 and max_target seqs=500`. We retrieved the full homologous sequence set (designated *HmgS1*) using their parsed sequence accessions and then performed overlap reduction (using the 30% threshold applied at the initial homology reduction process) against the *combined validation and testing set (* CVTS); using *HmgS1* as the query. Subsequently, we used the proportion of CVTS sequences overlapping with the *HmgS1* as a proxy for the extent of overlap. That is, we divided the number of overlapping sequences by the total number of sequences within CVTS. The rationale is that using overlapping sequences to generate profiles propagates the overlap (leakage) to the resulting sequence encoding.

#### Ablation of augmenting sequences from homology reduction in leakage calculation

To nullify the contribution of the overlapping sequences isolated within the training set to leakage calculation, we created *random-training-seqs-B* by removing the augmenting sequences resulting from the homology reduction process (shown as broken arrows in **Fig 2**) from *random-training-seqs-A*. A total of 384 sequences were removed, with 896 remaining (*random-training-seqs-B*), which constituted 10% of the corresponding training set (the single-localising subset of *TrnS5* with all augmenting sequences removed). We then pruned *HmgS1* of all hits associated with the 384 sequences. Subsequently, we performed overlap reduction on the test set using the remaining portion – *HmgS2*. This allowed us to observe the impact of these specific sequences on the extent of overlap – net leakage.

### Dataset benchmarking

To benchmark the reliability of our newly developed SCL dataset (*SCL2205*; n = 19,074), we assessed its performance against a comparable SoTA: DeepLoc2, redundancy-reduced at an 80% threshold (SwissProt train-validation (*DEEP-TV*); n = 22,126). Finally, we evaluated these using two independent sets: SwissProt sorting signal (*DEEP-SS*) and human protein atlas (*DEEP-HPA*) [29].

#### Dataset pre-processing

*SCL2205* and *DEEP-TV* were redundancy-reduced similarly. For training, *SCL2205* and *DEEP-TV* were respectively split into *training* (*n* = 10,680; 12,390, 56%), *validation* (*n* = 2,671; 3,098 [14%]) and *testing* (*n* = 5,723; 6,638, 30%) partitions. The splitting was random and stratified on class labels. All *DEEP-TV* sequences localising to the Golgi apparatus were excluded because they had fewer than *n* = 100 counts, which was the per-class count threshold used with *SCL2205*. For brevity, experiments were performed using only the single-label classes, which were intuitively coded *SCL2205*_*s*_ and *DEEP-TV*_*s*_, respectively. See (**Table 2**) for the class distribution across datasets and partitions.

**Table 2.** Label-wise distribution across datasets and partitions (Train [Trn], Validation [Vld], and Testing [Tst]) for single-location labels only. There is a notable difference for the *Membrane* class, resulting from the divergent binning approach adopted in the datasets. *DEEP-TV* has a higher number of training examples than *SCL2205* in 60% of the classes. For brevity: Membrane (MEM), Nucleus (NUC), Secreted (SEC), Cytoplasm (CYT), Mitochondrion (MIT), Plastid (PLA), Endoplasmic reticulum (ER), Cell projection (CEP), Peroxisome (PER) and Lysosome/vacuole (LYS/VAC).

| Label | *SCL2205<sub>s</sub>* |  |  | *DEEP-TV<sub>s</sub>* |  |  |
| --- | --- | --- | --- | --- | --- | --- |
|  | Trn | Vld | Tst | Trn | Vld | Tst |
| MEM | 2642 | 661 | 1415 | 1117 | 599 | 279 |
| NUC | 2340 | 585 | 1254 | 2581 | 1383 | 646 |
| SEC | 1475 | 369 | 791 | 1307 | 701 | 327 |
| CYT | 1380 | 345 | 740 | 2073 | 1111 | 518 |
| MIT | 530 | 132 | 284 | 617 | 331 | 154 |
| PLA | 329 | 82 | 176 | 332 | 178 | 83 |
| ER | 104 | 26 | 55 | 131 | 70 | 33 |
| CEP<sup>†</sup> | 98 | 25 | 53 | NA | NA | NA |
| PER | 76 | 19 | 41 | 75 | 40 | 19 |
| LYS/VAC<sup>†</sup> | NA | NA | NA | 106 | 57 | 26 |
| **Totals** | **8974** | **2244** | **4809** | **8339** | **4470** | **2085** |
| **Coverage** | **56%** | **14%** | **30%** | **56%** | **14%** | **30%** |

<sup>†</sup> Missing for the corresponding dataset and omitted during dataset-comparison model training.

Since the *DEEP-SS* independent dataset was initially labelled by sorting signals, it was therefore relabelled. We retrieved subcellular location labels from UniProtKB using persistent sequence accessions. Only sequences with matching UniProtKB accessions were considered. The human protein atlas (HPA) dataset had labels corresponding to our training data.

#### Test modelling

##### Input representation

Representation was either tokenizer- or protein language model (PLM)-based, depending on the testing model architecture – CNN or PLM embedding network, and a common fully connected neural network (FCNN) classification network.

##### Architecture

While the *ad hoc* CNN architecture is briefly described in **Supplementary File S1**, for PLM embedding, we used an independent PLM – *Rostlab/prot t5 xl uniref50* – by the Rost Lab [9], using the *Transformers* Application Programming Interface (API) [32] from Hugging Face.

##### Training

Training was run until the model overfitted. That is, a sustained increase in (training-validation) loss after an initial decrease, as observed in the loss plot. For computational robustness (training was on a managed computing cluster), each model training phase was run three times, followed by weight averaging to instantiate the final model used for testing. The model snapshot with the best validation accuracy just before the sustained loss increase was saved for each phase.

##### Evaluation

The test evaluations were compared using multiple strategies, in isolation or as combinations: *macro* and *per-class* PR-AUC metrics for ranking and positive retrievals, McNemar’s error rates test, and stratified bootstrapping on metrics differences uncertainty quantification. Evaluation testing was performed using independent test sets *DEEP-SS* and *DEEP-HPA*, see **Table 3**.

**Table 3.** Label-wise counts for independent test datasets.

| Label | *DEEP-SS* | *DEEP-HPA* |
| --- | --- | --- |
| Secreted<sup>†</sup> | 522 | NA |
| Membrane | 170 | 161 |
| Mitochondrion | 111 | 175 |
| Peroxisome | 77 | 7 |
| Nucleus | 63 | 664 |
| Cytoplasm | 10 | 263 |
| ER | 5 | 31 |
| Plastid<sup>†</sup> | 1 | NA |
| **Totals** | **958** | **1301** |

<sup>†</sup> Missing for the corresponding dataset.

### SCL2205_*s*_ *versus* DEEP-TV_*s*_ *comparison*

To compare datasets as holistic stand-alone entities, we used the corresponding partitions of *SCL2205*_*s*_ and *DEEP-TV*_*s*_ to train *ad hoc* CNN-based model; we also isolated model effects by training an independent PLM-based model. The models resulting from training with *SCL2205*_*s*_ and *DEEP-TV*_*s*_; which for brevity we will, respectively, call *Model A* and *Model B*, were then evaluated. The independent test datasets were overlap-reduced with each corresponding *training* set, see details in **Supplementary File S1** for the resulting ablation counts.

### DEEP-SS and **DEEP-HPA** independent test sets

We highlight distinct characteristics of these two external evaluation sets. *DEEP-SS* is drawn from universal protein (UniProt)-SwissProt, which is also the source of the training datasets; therefore, it can be considered an *in-distribution* set. In contrast, *DEEP-HPA* is based on SCL annotations from the HPA, which employs a different annotation strategy. SCL in HPA is diversely investigated; using isolated or integrated techniques comprising of immunocytochemistry-immunofluorescence (ICC-IF), confocal microscopy, and staining [28]. Moreover, the distinction is highlighted by the extent of overlap with the *SCL2205* and *DEEP-TV* training sets, as captured in various overlap-reduction instances, see **Tables 2 and 3** in **Supplementary File S1**. Furthermore, whereas the UniProt-SwissProt data covers multiple taxa, HPA covers only humans. Therefore, we consider *DEEP-HPA* as an *out-of-distribution* (OOD) set.

## Results

### Mapping improved generalisation

We employed manual label mapping to increase target diversity and boost the number of training samples, to improve model generalisability and performance. A representative outcome of the mapping process using *Plastid* – the most impacted location – is illustrated in **Fig 3**. For clarity, we refer to the model using label mapping as *Model A* and the model using native labelling as *Model B*.

**Fig. 3.**
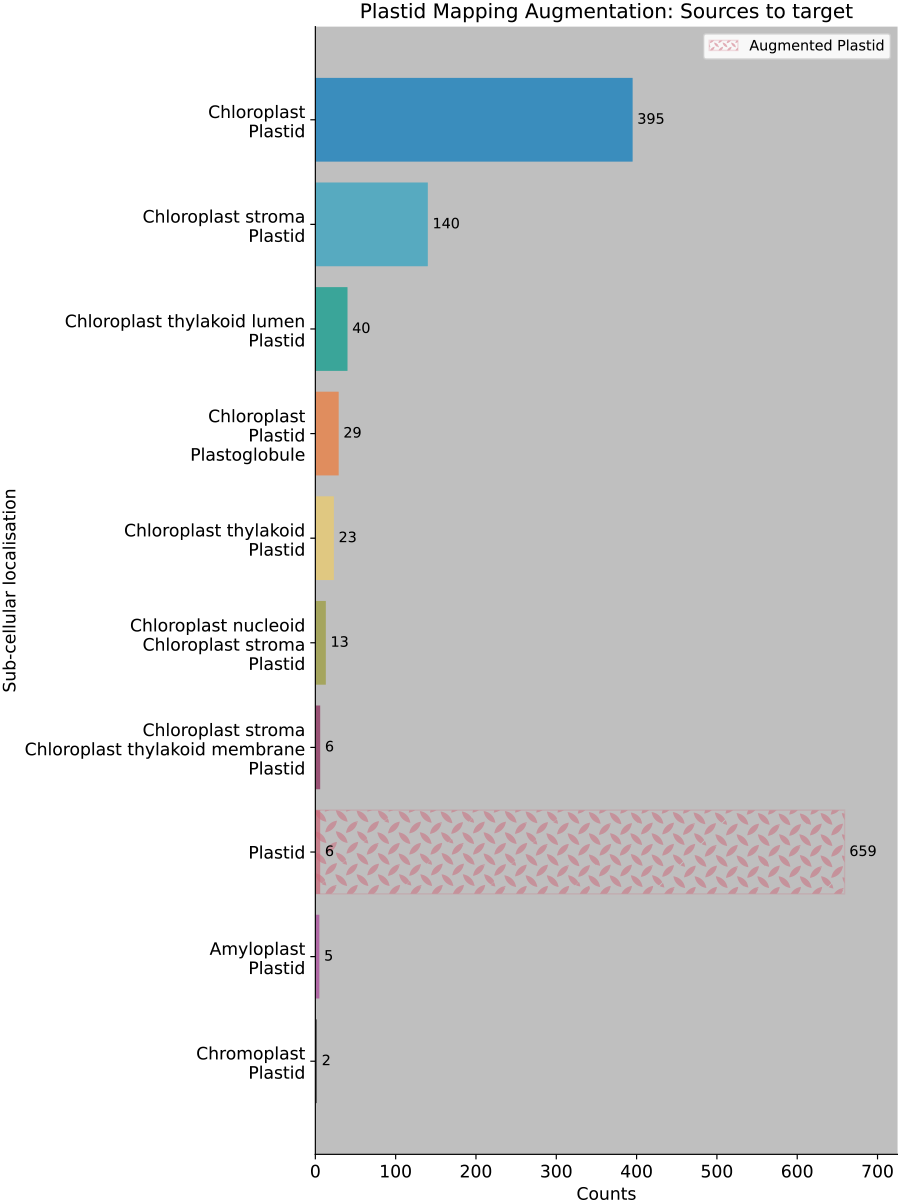
Frequencies for the Plastid label mapping augmentation involving nine locations. We reported the highest fold increase for Plastid (103.67 fold). **Alt text**: Graphics of a mapping case study involving Plastid, illustrating a fold change of over 100.

We observed a substantial improvement of the macro PR-AUC for *DEEP-SS* when using label mapping (an increase of 9.0% points); however, the improvement for *DEEP-HPA* was marginal (1.5% points). Full details, including per-class PR-AUC with their respective prevalences, are provided in the **Results** section of **Supplementary File S1**. Overall, label mapping enhanced model performance across most classes; however, we observed a slight deterioration in the *Cytoplasm* and *Secreted* categories for *DEEP-SS*, and the *Nucleus* category for *DEEP-HPA*.

An asymptotic McNemar’s test with continuity correction revealed a significant performance difference between *Model A* and *Model B* on both test sets. This test evaluates error rates at a fixed threshold. For *DEEP-SS, χ*^2^(*df* = 1) = 8.2, *p* < 0.004, with *Model A* outperforming *Model B* (b = 108, c = 69; *Effect Size Z* = 2.9). In contrast, for *DEEP-HPA, χ*^2^(*df* = 1) = 278.8, *p* < 0.001, with *Model B* outperforming *Model A* (*b* = 610, *c* = 149; *Effect Size Z* = −16.7).

The significant performance difference between *Model A* and *Model B* was supported by the stratified paired bootstrap confidence interval (CI) test. However, whereas *Model A* still, as in the McNemar’s test, outperformed *Model B* on the *DEEP-SS* set, the reverse was observed for the *DEEP-HPA* set in this test, where the label mapping of *Model A* proved superior. The stratified paired bootstrap CI test evaluates the ranking and retrieval of positives. **Fig 4** illustrates the uncertainty levels based on the macro PR-AUC differences between the two models.

**Fig. 4.**
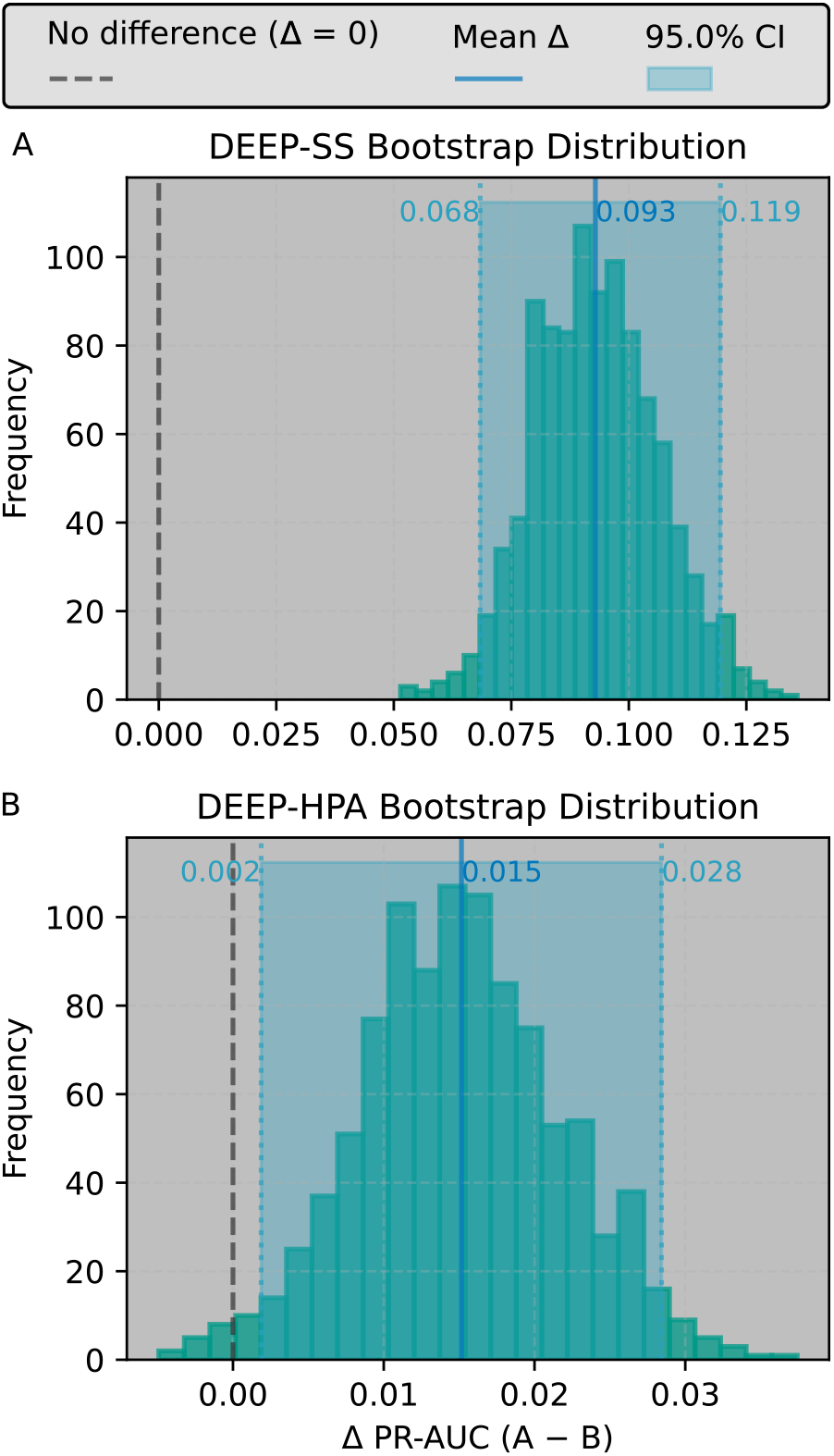
Histogram of the distribution of 1000 bootstrap PR-AUC-metric differences between models *A* and *B* as trained on 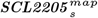 and 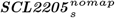, respectively. The top panel (**A**) was tested on *DEEP-SS* (*A* > *B*), whereas the bottom panel (**B**) was tested on *DEEP-HPA* (*A* > *B*). **Alt text**: Graphics of a histogram illustrating the effect of label mapping using two independent test datasets. Panel A (top) represents the in-distribution UniProtKB test data, while Panel B (bottom) represent the out-of-distribution human protein atlas test data.

### Data leakage from homology augmentation

We show and quantify that homology augmentation can partially undo prior *training-testing* dataset homology reduction. Interestingly, we found that the 10% training set sample resulted in 4.8% *training-testing* sequence overlap. *HmgS1* had 664,719 sequences, while *HmgS2* had 477,870, a 28.1% (*n* = 186, 849) reduction of the initial homology-augmenting sequence hits. Consequently, there was a 22.9% decrease in the number of sequences overlapping the test set between using *HmgS1* (*n* = 201) and *HmgS2* (*n* = 155) as query. These results underscore the need for careful consideration when implementing homology search augmentation of training data, which may inflate performance metrics and/or hamper generalisation. They also provide a baseline quantification, arguably the first such indication, for the amount of overlap resulting from database search augmentation, on the same similarity measurement scale.

### A benchmark dataset for subcellular localisation prediction

We developed *SCL2205* from the latest UniProtKB release (UniProtKB/Swiss-Prot: Release 2022 05; 20230124), with extensive quality preprocessing to ensure reliability, manual-mapping augmentation to increase training data and improve generalisation, and stringent data partitioning to minimise data leakage. While we observed a 1.71-fold increase in data during the development process, the inherent imbalance across data classes persists, albeit reduced. Therefore, we opted for the macro PR-AUC metrics to evaluate performance relative to other datasets. The precision–recall curve (PRC) curve evaluates recall (sensitivity/true positive rate (TPR)) versus precision (specificity/positive predictive value (PPV)) at multiple thresholds to highlight the trade-off between them. On the other hand, the PR-AUC score summarises the PRC curve to a number. Unlike in area under the ROC curve (ROC-AUC), where the binary baseline is 0.5, positive-class prevalence is the baseline in PR-AUC.

Next, we consider the results for the two model architectures used for dataset comparisons. **Table 4** provides the global PR-AUC scores, while the per-class scores are summarised in **Supplementary File S1**.

**Table 4.** *SCL2205* _*s*_ and *DEEP-TV* _*s*_ performance comparisons: a summary of the macro PR-AUC across model variants and test datasets. Despite using vanilla models, all scores exceeded the expected random guessing level. We observed the most significant gains in the PLM-based model – up to 10.8% points in favour of *SCL2205* _*s*_.

| Dataset & Architecture | Baseline Prevalence | *SCL2205<sub>s</sub>* 8-class | *DEEP-TV<sub>s</sub>* 8-class |
| --- | --- | --- | --- |
| *DEEP-SS<sub>CNN</sub>* | 0.143* | 0.257 | 0.334 |
| *DEEP-HPA<sub>CNN</sub>* | 0.167* | 0.194 | 0.184 |
| *DEEP-SS<sub>PLM</sub>* | 0.143* | 0.561 | 0.453 |
| *DEEP-HPA<sub>PLM</sub>* | 0.167* | 0.345 | 0.359 |

\* We computed the average prevalence across classes for each test set.

#### CNN-based architecture

An asymptotic McNemar’s test with continuity correction showed a significant difference between the models trained on *SCL2205* _*s*_ (*Model A*) and *DEEP-TV* _*s*_ (*Model B*) when evaluated on the two test sets. For *DEEP-SS, χ*^2^(*df* = 1) = 18.1, *p* < 0.001; (*B* > *A* [*b* = 90, *c* = 158]) with an Effect Size *Z* = −4.3. *DEEP-HPA*: *χ*^2^(*df* = 1) = 143.0, *p* < 0.001; (*A* > *B* [*b* = 145, *c* = 0]) with an Effect Size *Z* = 12.0. We observed a similar pattern using the stratified bootstrap on the PR-AUC metric difference test (Δ = *A* − *B*). For *DEEP-SS*, 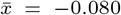, *p* < 0.001; (*B* > *A* [95% *CI* : −0.136 − −0.043]), see **Fig 5, Panel A**. However, although *Model A* exhibits a higher PR-AUC than *Model B*, the difference was not significant for *DEEP-HPA*: 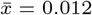, *p* < 0.094; (*Unclear direction* [95% *CI* : −0.001 − 0.031]), see **Fig 5, Panel B**.

**Fig. 5.**
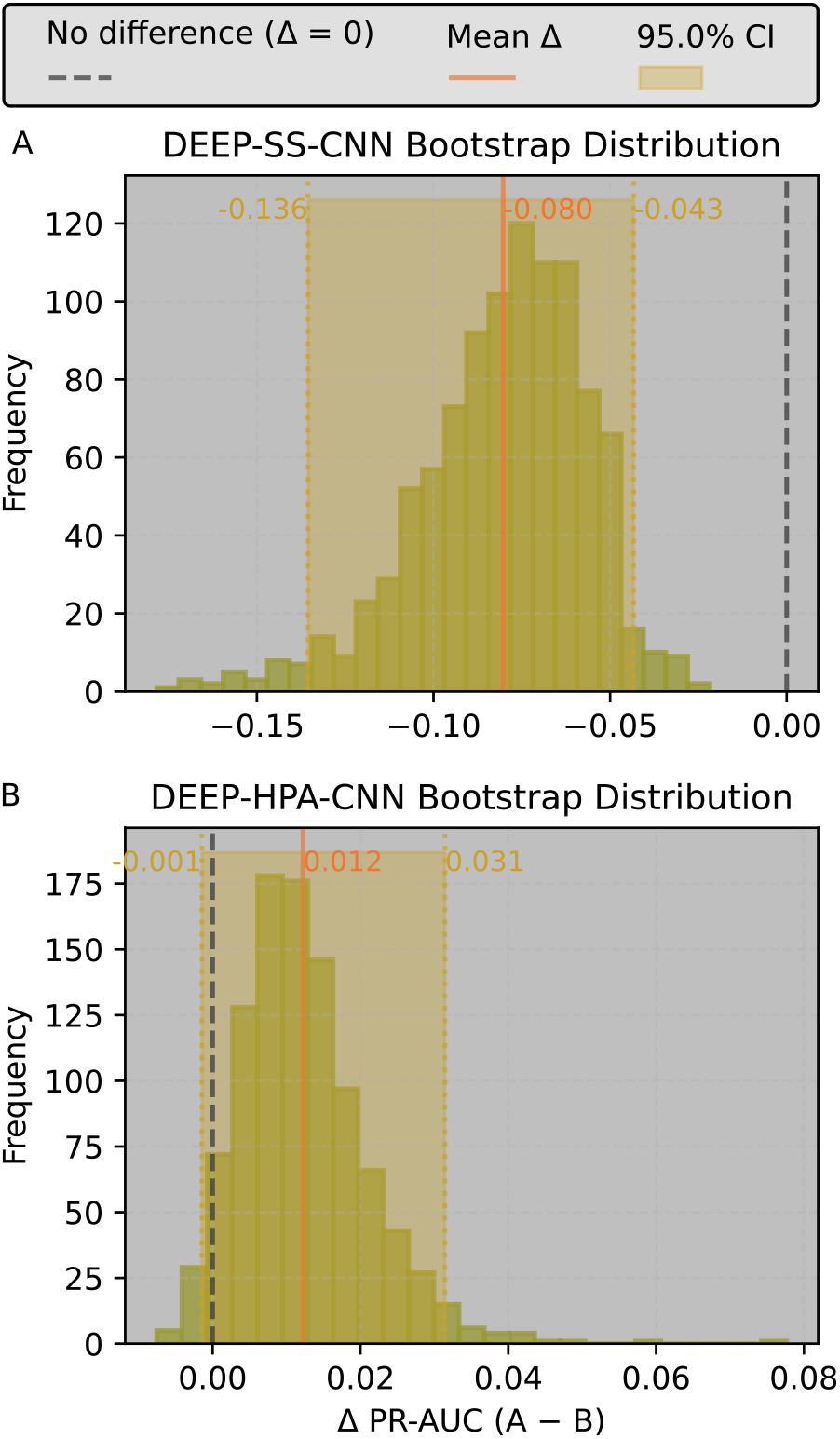
Histogram of the distribution of 1,000 bootstrap PR-AUC-metric differences between CNN-based models *A* and *B* as trained on *SCL2205* _*s*_ and *DEEP-TV* _*s*_, respectively. The top panel (**A**) was tested on *DEEP-SS* (*B* > *A*), whereas the bottom panel (**B**) was tested on *DEEP-HPA* (*No significant difference*). **Alt text**: Graphics of a histogram illustrating the performance difference between our dataset and a SoTA based on a CNN model architecture. Two independent test datasets were used, an in-distribution UniProtKB dataset (Panel A [top]) and an out-of-distribution human protein atlas dataset (Panel B [bottom]).

#### PLM-based architecture

Similar to the above architecture, we implemented tests on the PLM-based models. Interestingly here, we observed the reverse trend for *DEEP-SS* : *χ*^2^(*df* = 1) = 67.0, *p* < 0.001; (*A* > *B* [*b* = 90, *c* = 8]) with an Effect Size *Z* = 8.3, while the difference was not significant for *DEEP-HPA*: *χ*^2^(*df* = 1) = 3.5, *p* < 0.060; (*Unclear direction* [*b* = 109, *c* = 82]) with an Effect Size *Z* = 2.0.

In contrast, for the stratified bootstrap on the PR-AUC metric difference test (Δ = *A* − *B*) for *DEEP-SS* : 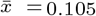, *p* < 0.001; (*A* > *B* [ 95% *CI* : 0.084 − 0.125]), see **Fig 6, Panel A**. Nonetheless, despite *Model B* depicting a higher PR-AUC than *Model A*, the difference was not significant, just as before; *DEEP-HPA*: 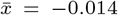, *p* < 0.07; (*Unclear direction* [95% *CI* : −0.029 − 0.001]), see **Fig 6, Panel B**.

**Fig. 6.**
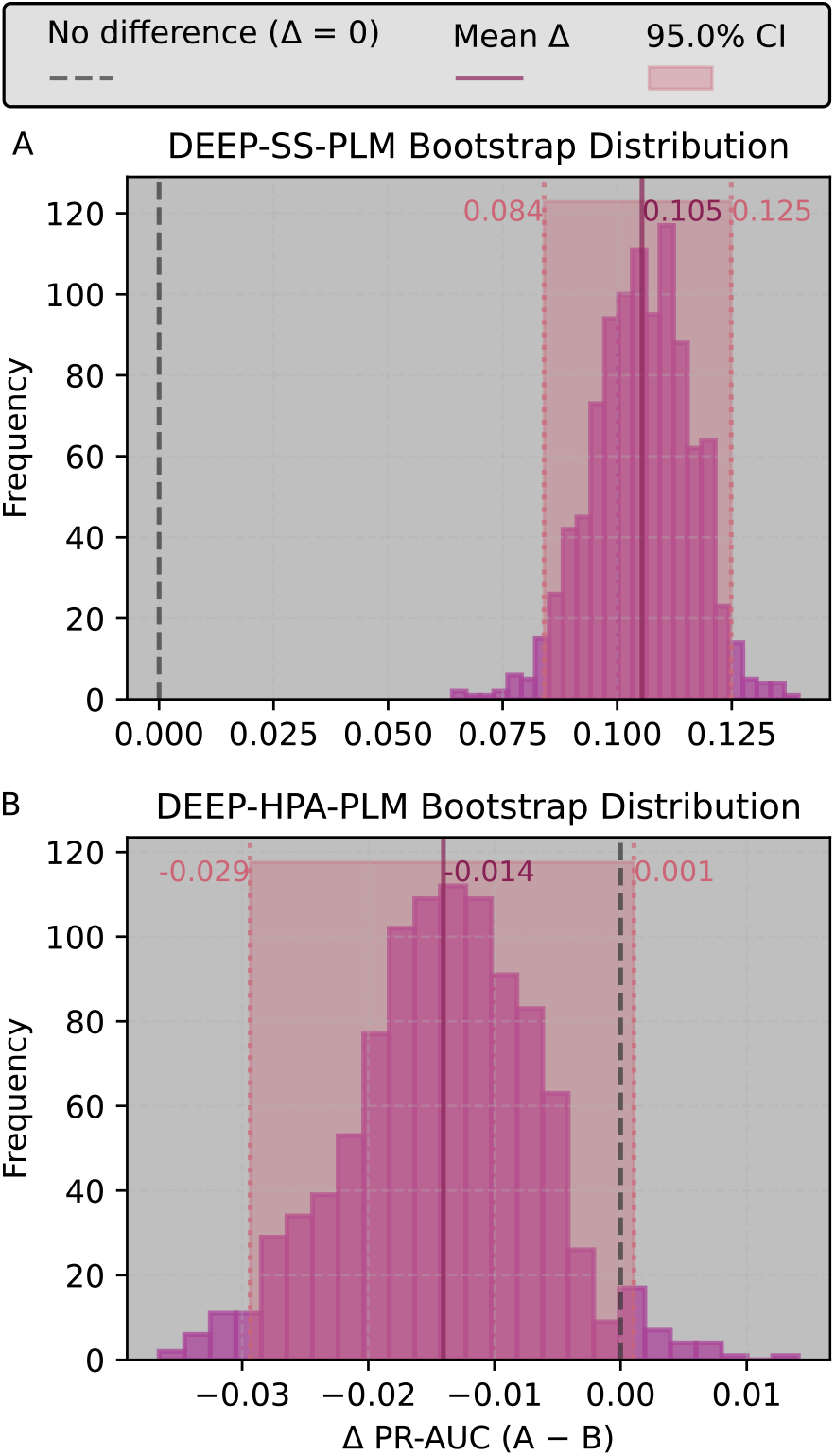
Histogram of the distribution of 1,000 bootstrap PR-AUC-metric differences between PLM-based models *A* and *B* as trained on *SCL2205* _*s*_ and *DEEP-TV* _*s*_, respectively. The top panel (**A**) was tested on *DEEP-SS* (*A* > *B*), whereas the bottom panel (**B**) was tested on *DEEP-HPA* (*No significant difference*). **Alt text**: Graphics of a histogram illustrating the performance difference between our dataset and a SoTA based on a PLM model architecture. Two independent test datasets were used, an in-distribution UniProtKB dataset (Panel A [top]) and an out-of-distribution human protein atlas dataset (Panel B [bottom]).

Overall, the choice of model architecture influenced the performance of the training data. The PLM-based models had lower uncertainty compared to CNN-based models; **Fig 6** and **Fig 5**.

### The final SCL2205 dataset

We propose *SCL2205*, a “leak-proof” dataset for SCL modelling that incorporates expert manual curation. This approach ensures high-quality training input, thereby favouring efficient model development. The specific compositions for the two tracks of the *SCL2205* dataset (*n* = 19074) are provided in the **Results** section of **Supplementary File S1**. Furthermore, we provide a convenient, open-source interface (`p-scldata`) to facilitate easy access to the dataset. This is available through the public Python Package Index (PyPI), ensuring that *SCL2205* is easily integrated into existing bioinformatics workflows.

## Discussion

The rapid advancement of AI in genomics has created an urgent need for robust, standardised datasets. *SCL2205* addresses this gap by providing a meticulously curated resource for sequence-based spatial modelling. By implementing a rigorous homology-reduction procedure, we ensure that AI models are tested against truly independent data, preventing the ‘overfitting’ that often plagues genomic predictions. This work establishes a new benchmark for the PLM era, offering a transparent and scalable foundation for characterising the protein landscape with precision.

Inherently, DL modelling requires large volumes of training data. While true, studies have shown that smaller, carefully curated datasets can improve model performance [21]. Although pre-training data sources in NLP are comparatively less controlled than UniProtKB, the latter still contains poor-quality data that requires pruning.

In the life sciences, we are often confronted with life-critical problems that demand more stringent performance controls. Therefore, we employed a variety of filters and, importantly, careful manual curation, to ensure high-quality pre-training SCL data. For instance, unlike in [3] and [29], where the UniProtKB annotation quality scores are ignored, probably because they are associated with the overall sequence annotation (not SCL only), we include them in filtering. By considering annotation quality scores, we emphasise the protein’s functional interrelationships as inseparable biological processes. Nonetheless, it may also be argued that protein functional annotation is progressive and that certain aspects of a protein may be better understood than others at any given time. However, highly beneficial insights can be obtained from a broadly understood basis, as reflected in the overall annotation quality score, thereby promoting discovery. Therefore, the annotation score metric is an important and readily accessible quality control component in pre-training data development.

Another key technical distinction of the *SCL2205* dataset is our decision to retain a sequence cut-off of up to 5,000 amino acids, avoiding the common practice of aggressive truncation. While many SoTA predictors truncate sequences to 1,000 residues to reduce computational overhead, such choices risk discarding critical biological information. Our approach is premised on signal positional agnosticism; we assume that SCL signals – such as C-terminal *ER*-retention motifs (e.g., Lys-Asp-Glu-Leu (KDEL)) or internal nuclear localisation signals – can occur anywhere within a protein’s primary structure.

By preserving nearly the entire sequence length distribution of the original UniProtKB records, *SCL2205* ensures that bidirectional architectures, including bidirectional long-short-term memory (bi-LSTMs) and the latest PLMs, can leverage terminal information from both the *N* - and *C*-termini. Truncation at the 1,000-residue mark would effectively “blind” the backward pass of a bi-LSTMs or the global attention mechanism of a *Transformer* to the C-terminal context of larger proteins. Consequently, *SCL2205* provides a more biologically consistent representation, particularly for the notable fraction of the eukaryotic proteome that exceeds standard truncation limits, thereby mitigating the risk of false negatives for locations defined by non-amino-terminal signals.

SCL datasets are inherently imbalanced, which contributes to the omission of low-representation classes from modelling, as evidenced in DL predictors [17, 2]. With low target diversity, generalisation is bound to suffer. Therefore, although with some precision trade-off, we alleviated the problem by manually mapping sequences to their higher-order subcellular locations as was illustrated in (**Fig 1**).

Notably, from **Table 1**, the number of examples for certain locations – such as the Membrane – drastically increased, a category like Plastid would be excluded from modelling by, say, a mapping logic just considering the term “plastid”, which is broadly defined in UniProtKB’s SCL ontology.

To validate the mapping process, we use a bar chart (**Fig 3**) to illustrate the exact mapping for *Plastid* – the most enriched location – as a representative case. When considered alongside the individual definitions from **Gene Ontology and GO Annotations**, we can justify the observed logic. For example, a *Plastid* is defined as: *“Any member of a family of organelles found in the cytoplasm of plants and some protists, which are membrane-bounded and contain DNA*.*”*

However, nuanced cases sometimes required a “voting” strategy. For instance, in the case of *Chloroplast stroma;Chloroplast thylakoid membrane;Plastid*, the majority of annotations favoured a generalised mapping to *Plastid* instead of *Membrane*. In instances where a “draw” occurred between two locations of interest, such as *Chloroplast thylakoid membrane;Plastid*, we argued that mapping to *Membrane* provided a more definitive biological classification. While it is alternatively arguable that mapping to *Plastid* supports a broader generalisation, these rare edge cases represent the inherent trade-offs in mapping-based data augmentation.

Consequently, manual intervention, though labour-intensive, not only increased the number of training examples but also improved class diversity. As our results demonstrate, this directly benefits model performance. Although location imbalance persists, it has been notably reduced. Therefore, *SCL2205* provides a robust foundation for further augmentation mechanisms, particularly during model development. We are addressing this in ongoing research and invite the community to explore these possibilities using the *SCL2205* dataset.

Our label-mapping results suggest a compelling enhancement to model ranking quality, especially for the PLM-based model, while underscoring the complexity of the quality-versus-quantity trade-off in spatial modelling. The two statistical tests employed, McNemar’s test and stratified bootstrapping CI on PR-AUC differences, address different notions of “superiority”. While the PR-AUC comparison determines which model better ranks and retrieves positives across various thresholds (quality), McNemar’s test evaluates which model performs better at a fixed decision threshold (the decision rule).

In the case of *DEEP-SS*, the significant performance boost suggests that broader, mapped labels helped the model identify general biological “rules” for sorting signals. However, the results for *DEEP-HPA* tell a conflicting story across the tests. Here, the native labelling (*Model B*) outperformed the mapped version on the McNemar’s test, whilst the opposite was observed with the bootstrapping test, indicating that improved ranking performance did not translate into superior hard classification performance at the chosen operating point. It also suggests that the high-precision human protein atlas data carry vital information that mapping or taxonomic heterogeneity may inadvertently “blur”.

We also evaluated a current SoTA augmentation approach, homology augmentation (see **Paragraph 2.1** for definition), to assess its impact on data partitioning. Previous studies have demonstrated the benefits of incorporating evolutionary information into both structure prediction [4, 16] and SCL prediction [24, 1]. These benefits may stem from improved generalisation, as shown by Nair et al., [24], who observed that SCL could be transferred with up to 90% accuracy for proteins sharing as little as 50% sequence identity.

Interestingly, their findings also indicated that SCL transfer remains feasible even at lower identity thresholds, similar to observations in structure prediction [23], particularly for major cellular compartments, which are overrepresented in databases. Despite these advantages, however, certain issues are either perpetuated or newly introduced. While class imbalance is inherent, data leakage remains a critical concern, especially in SoTA scenarios that claim “stringent train–test data partitioning”. We conjectured residual leakage due to homology augmentation, especially for classification, where sequence labels are inherent despite using unlabelled sequences for augmentation. While testing this hypothesis may require a more complex study, primarily due to the intensive resource requirements for homology searches, we tested a simpler proxy. Homology augmentation is a standard practice in sequence-based DL SCL predictors. It is anchored on conserving evolutionary function and has been shown to improve prediction accuracy relative to *sequence-feature-based-only* approaches [24, 12]. Although it can amplify signals using profiles of homologous proteins, common implementations have been shown to bias model evaluation [30]. The bias results from the overlap between the training and testing datasets.

We demonstrated that data leakage of at least 4.8% occurred, based on the same overlap-reduction (partitioning) approach applied prior to homology augmentation, when only 10% of the examples in the training set were used for homology augmentation. In contrast, previous efforts to examine biases in comparing profile- and non-profile-based SCL predictors, [30] used an *ad hoc* similarity proxy that differed from the partitioning methods of the models under consideration – potentially obscuring the true extent of overlap.

Theoretically, our findings suggest that achieving 100% data leakage would require fewer examples than the full training set; however, this would depend on factors such as the degree of homology in the original dataset and the extent of overlap between the training/testing datasets, and the homology augmentation database. Regardless, we hypothesise that, relative to common practices in homology augmentation, such as MSA and position-specific scoring matrices, the extent of data leakage would be even greater than that observed with our 10% subset of native sequence data. While our similarity comparisons are conducted on a native-to-native, sequence-to-sequence basis, MSA- and PSSM-based approaches typically compare an averaged (consensus) sequence against a native sequence. Because a consensus sequence is intrinsically more likely to produce a broader range of hits, this implies greater overlap and, consequently, increased leakage, potentially even across different subcellular locations.

Therefore, at a minimum, predictors applying homology reduction followed by homology augmentation must assess and report any post-homology-augmentation training–testing overlap. We acknowledge that rigorously evaluating this is challenging, as the sequence representation following augmentation differs from that used during initial reduction unless equivalent consensus sequences are employed.

In light of the above, pursuing emerging alternative augmentation strategies, [14, 25, 9, 29], may be a superior option. Interestingly, we observed that *SCL2205* better complements the new frontier of PLM by exhibiting improved generalisation over *DEEP-TV* for the *in-distribution* set (*DEEP-SS*), while exhibiting reduced performance on the OOD counter-set (*DEEP-HPA*; **Figure 6**). To delineate true effect from noise, we performed a heterogeneity check using Cochran’s *Q* test. The significant heterogeneity observed across the models suggests that the CNN and PLM architectures are learning fundamentally different signals from the same data.

Conversely, we observed contrasting results between *SCL2205* and *DEEP-TV* on both independent test sets when using the CNN-based model (textbfFigure 5). These outcomes can be attributed to several factors beyond architectural design. Foremost, *DEEP-HPA* is a unique OOD dataset due to its taxonomic bias; we therefore expect our multi-taxa dataset to notably diverge from it relative to *DEEP-TV*. Secondly, the training paradigm for PLMs aligns more naturally with our label-mapping approach. Thirdly, the statistical tests are tailored for different notions of “superiority”. Finally, differences in model calibration may contribute to overconfidence, particularly in PLMs [13, 8, 6]. However, both models were tested in their *vanilla* forms, without calibration or optimisation, and reliability diagrams depicted very similar profiles across the model variants.

These divergences highlight a critical trade-off in spatial modelling: the delicate balance between breadth (e.g., through mapping and/or taxonomic diversity) and depth (the precision of native labels and/or taxonomic specificity).

Overall, *SCL2205* alignsmore effectively than *DEEP-TV* with the increasing adoption of pre-trained PLMs in genomics. This distinction is critical for the PLM frontier: if a researcher requires a definitive “yes/no” prediction, the native labelling of *Model B* is preferable. Conversely, for large-scale genomic screening where ranking candidates is the priority, the augmented mapping of *Model A* provides greater utility.

Ultimately, we have proposed a rigorously developed dataset that enhances the trustworthiness of AI models. It facilitates efficient and environmentally sustainable development by mitigating “noisy” overhead data, whilst enriching generalisation through diversity. By providing *SCL2205* as an installable package (`p-scldata`), we adhere to open-science principles, ensuring seamless integration with modern ML frameworks and providing a compelling benchmark for future genomic discovery.

## Conclusion

The computational annotation of proteins with SCL remains a critical component of functional genome annotation. As the field enters new frontiers, such as PLMs, new challenges emerge – most notably regarding the quality and diversity of input data. While data are the primary driving force behind the AI revolution, acquiring high-quality datasets for training biological models remains a significant hurdle. Nevertheless, techniques such as data augmentation and human-aided curation have begun to alleviate these issues. In this study, we present *SCL2205* : a new, high-quality dataset that highlights the limitations of current state-of-the-art approaches to data development. Through independent validation, we demonstrate that *SCL2205* outperforms existing alternatives, particularly for applications involving PLM-based modelling. We acknowledge that certain persistent challenges, such as class imbalance, must still be addressed through further research beyond data preparation. Nonetheless, *SCL2205* serves as a trustworthy and reliable benchmark for AI application in SCL prediction, laying the foundation for future discoveries in the genomic landscape. Ultimately, by providing a more accurate spatial map of localisation within the cell, we move closer to a future where we can rapidly identify the molecular drivers of rare diseases and accelerate the development of life-changing targeted therapies.

## Data availability

Data are available through the following open-source avenues:

1. **DRYAD**: An archive (DOI) under the Creative Commons Zero (CC0 1.0) licence.
2. **PyPI**: An installable Python data package (p-scldata) under the MIT licence.
3. **Data & Code**: [GitHub](https://github.com/ousodaniel/scldata/blob/main)

## Author contributions statement

DO and GP conceived the experiments, DO conducted the experiments, DO analysed the results, and DO wrote the manuscript. DO and GP reviewed and edited the manuscript. GP supervised the study. The project funding was through GP.

## Acknowledgments

The authors thank the anonymous reviewers for their valuable suggestions. This research was funded by Research Ireland through the Centre for Research Training in Genomics Data Science under Grant number #18/CRT/6214.

