## Supplementary File S2 for "Developing *SCL2205*: A Protein Sequence-based Spatial Modelling Dataset for the Protein Language Model Frontier"

### Manual Subcellular Location Mapping and Harmonisation

1

Table 1: **Manual subcellular location mapping and harmonisation**. The columns comprise the mined original subcellular locations, their corresponding counts, and the mapped/harmonised annotations, respectively. The exercise was guided by the universal protein knowledgebase (UniProtKB)'s sub-cellular localisation (SCL) ontology information [1] and the interactive cell from *SwissBioPics* library [2].

| **Location** | **Counts** | **Mapped** |
|---|---|---|
| Secreted | 4877 | NA |
| Nucleus | 3695 | NA |
| Cytoplasm | 2876 | NA |
| Cytoplasm;Nucleus | 2350 | NA |
| Mitochondrion | 955 | NA |
| Membrane | 324 | NA |
| Cytoplasm;Cytoskeleton | 253 | NA |
| Endoplasmic reticulum | 150 | NA |
| Peroxisome | 145 | NA |
| Centrosome;Cytoplasm;Cytoskeleton;Microtubule organizing center | 96 | NA |
| Cytoplasm;Membrane | 92 | NA |
| Cell projection | 6 | NA |
| Plastid | 6 | NA |
| Cell membrane | 2310 | Membrane |
| Endoplasmic reticulum membrane | 888 | Membrane |
| Cell inner membrane | 602 | Membrane |
| Nucleolus;Nucleus | 486 | Nucleus |
| Mitochondrion inner membrane | 463 | Membrane |
| Chloroplast;Plastid | 395 | Plastid |
| Cytoplasm;Cytosol | 357 | Cytoplasm |
| Golgi apparatus membrane | 314 | Membrane |
| Chromosome;Nucleus | 313 | Nucleus |
| Cell wall;Secreted | 301 | Secreted |
| Mitochondrion matrix | 216 | Mitochondrion |
| Cell membrane;Cytoplasm | 200 | Cytoplasm;Membrane |
| Vacuole membrane | 193 | Membrane |
| Chloroplast thylakoid membrane;Plastid | 169 | Membrane |
| Cytoplasm;Cytosol;Nucleus | 158 | Cytoplasm;Nucleus |
| Chloroplast stroma;Plastid | 140 | Plastid |
| Cellular thylakoid membrane | 131 | Membrane |
| Mitochondrion outer membrane | 124 | Membrane |
| Cell outer membrane | 124 | Membrane |
| Extracellular matrix;Extracellular space;Secreted | 113 | Secreted |
| Cytoplasm;Nucleolus;Nucleus | 112 | Cytoplasm;Nucleus |
| Apical cell membrane | 112 | Membrane |
| Host nucleus | 109 | Nucleus |
| Mitochondrion membrane | 93 | Membrane |
| Extracellular space;Secreted | 87 | Secreted |
| Cytoplasm;Perinuclear region | 82 | Cytoplasm |
| Host cytoplasm | 81 | Cytoplasm |
| Nucleoplasm;Nucleus | 80 | Nucleus |
| Endoplasmic reticulum membrane;Golgi apparatus membrane | 78 | Membrane |

| Location | Counts | Mapped |
|---|---|---|
| Host cytoplasm;Host nucleus | 76 | Cytoplasm;Nucleus |
| Peroxisome membrane | 76 | Membrane |
| Nucleus speckle | 75 | Nucleus |
| Endoplasmic reticulum lumen | 72 | ER |
| Cell membrane;Endoplasmic reticulum membrane | 72 | Membrane |
| Cell projection;Cilium | 71 | Cell projection |
| Cell projection;Cilium;Flagellum | 68 | Cell projection |
| Lysosome membrane | 66 | Membrane |
| Centriole;Centrosome;Cytoplasm;Cytoskeleton;Microtubule organizing center | 64 | Centrosome;Cytoplasm;Cytoskeleton;Microtubule organizing center |
| Basolateral cell membrane | 56 | Membrane |
| Cell membrane;Membrane raft | 55 | Membrane |
| Nucleus;Nucleus speckle | 51 | Nucleus |
| Endosome membrane | 49 | Membrane |
| Cell cortex;Cytoplasm | 46 | Cytoplasm |
| Apical cell membrane;Cell membrane | 46 | Membrane |
| Nucleus membrane | 44 | Membrane |
| Microsome membrane | 43 | Membrane |
| Cytoplasm;Nucleus;Perinuclear region | 42 | Cytoplasm;Nucleus |
| Cytoplasm;Nucleoplasm;Nucleus | 41 | Cytoplasm;Nucleus |
| Chloroplast thylakoid lumen;Plastid | 40 | Plastid |
| Nucleolus;Nucleoplasm;Nucleus | 37 | Nucleus |
| Nucleus envelope | 36 | Membrane |
| Cytoplasm;Cytoskeleton;Microtubule organizing center;Spindle pole body | 34 | Centrosome;Cytoplasm;Cytoskeleton;Microtubule organizing center |
| Basolateral cell membrane;Cell membrane | 34 | Membrane |
| Nucleus inner membrane | 32 | Membrane |
| Endoplasmic reticulum membrane;Microsome membrane | 32 | Membrane |
| Endoplasmic reticulum membrane;Nucleus membrane | 29 | Membrane |
| Chloroplast;Plastid;Plastoglobule | 29 | Plastid |
| Nucleoplasm;Nucleus;Nucleus speckle | 28 | Nucleus |
| Cell membrane;Golgi apparatus membrane | 26 | Membrane |
| Cytoplasmic vesicle membrane | 26 | Membrane |
| Cell projection;Dendrite | 25 | Cell projection |
| Centrosome;Cytoplasm;Cytoskeleton;Microtubule organizing center;Nucleus | 25 | Centrosome;Cytoplasm;Cytoskeleton;Microtubule organizing center |
| Cell membrane;Postsynaptic cell membrane | 25 | Membrane |
| Cell membrane;Vacuole membrane | 25 | Membrane |
| Host cell membrane | 25 | Membrane |
| Centrosome;Cytoplasm;Cytoskeleton;Microtubule organizing center;Spindle | 24 | Centrosome;Cytoplasm;Cytoskeleton;Microtubule organizing center |
| Early endosome membrane | 24 | Membrane |
| Cell membrane;Cytoplasmic vesicle membrane | 24 | Membrane |
| Centriole;Centrosome;Cilium basal body; Cytoplasm;Cytoskeleton;Microtubule organizing center | 23 | Centrosome;Cytoplasm;Cytoskeleton;Microtubule organizing center |
| Chloroplast outer membrane;Plastid | 23 | Membrane |
| Cell membrane;Endosome membrane | 23 | Membrane |
| Preautophagosomal structure membrane | 23 | Membrane |
| Chloroplast thylakoid;Plastid | 23 | Plastid |

| Location | Counts | Mapped |
|---|---|---|
| Axon;Cell projection | 22 | Cell projection |
| Cytoplasm;Nucleolus;Nucleoplasm;Nucleus | 22 | Cytoplasm;Nucleus |
| Late endosome membrane;Lysosome membrane | 22 | Membrane |
| Endomembrane system | 22 | Membrane |
| Nucleus;Pml body | 22 | Nucleus |
| Cytoplasm;Nucleus;P-body | 20 | Cytoplasm;Nucleus |
| Cytoplasm;Cytosol;Membrane | 19 | Cytoplasm;Membrane |
| Late endosome membrane | 19 | Membrane |
| Host endoplasmic reticulum membrane | 16 | Membrane |
| Cellular chromatophore membrane | 16 | Membrane |
| Nucleus matrix | 16 | Nucleus |
| Peroxisome matrix | 16 | Peroxisome |
| Sarcoplasmic reticulum membrane | 15 | Membrane |
| Cell outer membrane;Cell surface | 15 | Membrane |
| Cell projection;Stereocilium | 14 | Cell projection |
| Cytoplasm;Cytoskeleton;Microtubule organizing center;Nucleus;Spindle pole body | 14 | Centrosome;Cytoplasm;Cytoskeleton;Microtubule organizing center |
| Centriolar satellite;Centrosome;Cytoplasm;Cytoskeleton; Microtubule organizing center | 13 | Centrosome;Cytoplasm;Cytoskeleton;Microtubule organizing center |
| Postsynaptic cell membrane | 13 | Membrane |
| Chloroplast nucleoid;Chloroplast stroma;Plastid | 13 | Plastid |
| Cell cortex;Cytoplasm;Nucleus | 12 | Cytoplasm;Nucleus |
| Apical cell membrane;Basolateral cell membrane | 12 | Membrane |
| Endoplasmic reticulum membrane;Vacuole membrane | 12 | Membrane |
| Centrosome;Cytoplasm;Cytoskeleton;Microtubule organizing center;Spindle pole | 11 | Centrosome;Cytoplasm;Cytoskeleton;Microtubule organizing center |
| Cytoplasm;Cytoplasmic granule | 11 | Cytoplasm |
| Host cytoplasm;Host perinuclear region | 11 | Cytoplasm |
| Cytoplasm;Cytoplasmic granule;Perinuclear region | 11 | Cytoplasm |
| Cell cortex;Cytoplasm;Cytoskeleton | 11 | Cytoplasm;Cytoskeleton |
| Cytoplasm;Cytoskeleton;Spindle;Spindle pole | 11 | Cytoplasm;Cytoskeleton |
| Cell membrane;Membrane | 11 | Membrane |
| Recycling endosome membrane | 11 | Membrane |
| Cajal body;Nucleus | 11 | Nucleus |
| Endoplasmic reticulum membrane;Mitochondrion membrane | 10 | Membrane |
| Membrane raft | 10 | Membrane |
| Cell membrane;Lysosome membrane | 10 | Membrane |
| Endosome membrane;Lysosome membrane | 10 | Membrane |
| Mitochondrion matrix;Mitochondrion nucleoid | 10 | Mitochondrion |
| Host nucleolus;Host nucleus | 10 | Nucleus |
| Cytoplasm;Nucleus;Pml body | 9 | Cytoplasm;Nucleus |
| Endosome membrane;Golgi apparatus membrane | 9 | Membrane |
| Centrosome;Cytoplasm;Cytoskeleton;Microtubule organizing center;Spindle;Spindle pole | 7 | Centrosome;Cytoplasm;Cytoskeleton;Microtubule organizing center |
| Cytoplasm;Nucleus;P-body;Stress granule | 7 | Cytoplasm;Nucleus |
| Cell membrane;Endosome membrane;Lysosome membrane | 7 | Membrane |
| Cytoplasm;Nucleoplasm;Nucleus;Nucleus speckle | 6 | Cytoplasm;Nucleus |
| Endoplasmic reticulum membrane;Membrane | 6 | Membrane |
| Cajal body;Nucleus;Nucleus speckle | 6 | Nucleus |

| **Location** | **Counts** | **Mapped** |
|---|---|---|
| Chloroplast stroma;Chloroplast thylakoid membrane;Plastid | 6 | Plastid |
| Centrosome;Cytoplasm;Cytoskeleton;Cytosol;Microtubule organizing center | 5 | Centrosome;Cytoplasm;Cytoskeleton;Microtubule organizing center |
| Centriolar satellite;Centriole;Centrosome; Cytoplasm;Cytoskeleton; Microtubule organizing center | 5 | Centrosome;Cytoplasm;Cytoskeleton;Microtubule organizing center |
| Centrosome;Cytoplasm;Cytoskeleton;Microtubule organizing center;Nucleus;Spindle | 5 | Centrosome;Cytoplasm;Cytoskeleton;Microtubule organizing center |
| Centrosome;Cilium basal body;Cytoplasm;Cytoskeleton; Microtubule organizing center | 5 | Centrosome;Cytoplasm;Cytoskeleton;Microtubule organizing center |
| Centromere;Centrosome;Chromosome;Cytoplasm; Cytoskeleton;Kinetochore;Microtubule organizing center;Spindle | 5 | Centrosome;Cytoplasm;Cytoskeleton;Microtubule organizing center |
| Cytoplasm;Cytosol;Nucleolus;Nucleus | 5 | Cytoplasm;Nucleus |
| Host endoplasmic reticulum | 5 | ER |
| Early endosome membrane;Recycling endosome membrane | 5 | Membrane |
| Golgi apparatus membrane;Vacuole membrane | 5 | Membrane |
| Endosome membrane;Vacuole membrane | 5 | Membrane |
| Amyloplast;Plastid | 5 | Plastid |
| Centrosome;Chromosome;Cytoplasm;Cytoskeleton; Microtubule organizing center;Nucleus | 4 | Centrosome;Cytoplasm;Cytoskeleton;Microtubule organizing center |
| Centriole;Centrosome;Cytoplasm;Cytoskeleton; Microtubule organizing center;Nucleus | 4 | Centrosome;Cytoplasm;Cytoskeleton;Microtubule organizing center |
| Cytoplasm;Cytoskeleton;Cytosol | 4 | Cytoplasm;Cytoskeleton |
| Nucleolus;Nucleoplasm;Nucleus;Nucleus speckle | 4 | Nucleus |
| Cytoplasm;Cytosol;Nucleoplasm;Nucleus | 3 | Cytoplasm;Nucleus |
| Cytoplasm;Nucleoplasm;Nucleus;Perinuclear region | 3 | Cytoplasm;Nucleus |
| Cell membrane;Multi-pass membrane protein | 3 | Membrane |
| Early endosome membrane;Golgi apparatus membrane | 3 | Membrane |
| Cell membrane;Recycling endosome membrane | 3 | Membrane |
| Cajal body;Nucleolus;Nucleus | 3 | Nucleus |
| Cytoplasmic vesicle;Extracellular space;Secreted;Secretory vesicle | 3 | Secreted |
| Centrosome;Cytoplasm;Cytoskeleton;Microtubule organizing center; Nucleolus;Nucleoplasm;Nucleus | 2 | Centrosome;Cytoplasm;Cytoskeleton;Microtubule organizing center |
| Host cytoplasm;Host cytosol | 2 | Cytoplasm |
| Cytoplasm;Nucleoplasm;Nucleus;Nucleus matrix | 2 | Cytoplasm;Nucleus |
| Cajal body;Cytoplasm;Nucleus | 2 | Cytoplasm;Nucleus |
| Cajal body;Cytoplasm;Nucleoplasm;Nucleus | 2 | Cytoplasm;Nucleus |
| Cajal body;Cytoplasm;Nucleus;Nucleus speckle | 2 | Cytoplasm;Nucleus |
| Cytoplasm;Nucleus;Nucleus speckle;P-body;Pml body | 2 | Cytoplasm;Nucleus |
| Cytoplasm;Nucleolus;Nucleoplasm;Nucleus;Nucleus speckle | 2 | Cytoplasm;Nucleus |
| Cytoplasm;Nucleoplasm;Nucleus;Pml body | 2 | Cytoplasm;Nucleus |
| Cytoplasm;Nucleus;Perikaryon | 2 | Cytoplasm;Nucleus |

| Location | Counts | Mapped |
| --- | --- | --- |
| Sarcoplasmic reticulum | 2 | ER |
| Cell membrane;Endomembrane system;Endosome membrane | 2 | Membrane |
| Glyoxysome membrane;Peroxisome membrane | 2 | Membrane |
| Cell membrane;Prospore membrane | 2 | Membrane |
| Apical cell membrane;Membrane | 2 | Membrane |
| Cell inner membrane;Cell outer membrane | 2 | Membrane |
| Nucleus envelope;Nucleus membrane | 2 | Membrane |
| Nucleoplasm;Nucleus;Pml body | 2 | Nucleus |
| Chromoplast;Plastid | 2 | Plastid |
| Chromosome;Cytoplasm;Nucleus;Pml body | 1 | Cytoplasm;Nucleus |
| Cytoplasm;Cytoplasmic vesicle;Nucleus;Pml body | 1 | Cytoplasm;Nucleus |

#### References

1. The UniProt Consortium, Alex Bateman, Maria-Jesus Martin, Sandra Orchard, Michele Magrane, Shadab Ahmad, Emanuele Alpi, Emily H Bowler-Barnett, Ramona Britto, Hema Bye-A-Jee, Austra Cukura, Paul Denny, Tunca Dogan, ThankGod Ebenezer, Jun Fan, Penelope Garmiri, Leonardo Jose da Costa Gonzales, Emma Hatton-Ellis, Abdulrahman Hussein, Alexandr Ignatchenko, Giuseppe Insana, Rizwan Ishtiaq, Vishal Joshi, Dushyanth Jyothi, Swaathi Kandasaamy, Antonia Lock, Aurelien Luciani, Marija Lugaric, Jie Luo, Yvonne Lussi, Alistair MacDougall, Fabio Madeira, Mahdi Mahmoudy, Alok Mishra, Katie Moulang, Andrew Nightingale, Sangya Pundir, Guoying Qi, Shriya Raj, Pedro Raposo, Daniel L Rice, Rabie Saidi, Rafael Santos, Elena Speretta, James Stephenson, Prabhat Totoo, Edward Turner, Nidhi Tyagi, Preethi Vasudev, Kate Warner, Xavier Watkins, Rossana Zaru, Hermann Zellner, Alan J Bridge, Lucila Aimo, Ghislaine Argoud-Puy, Andrea H Auchincloss, Kristian B Axelsen, Parit Bansal, Delphine Baratin, Teresa M Batista Neto, Marie-Claude Blatter, Jerven T Bolleman, Emmanuel Boutet, Lionel Breuza, Blanca Cabrera Gil, Cristina Casals-Casas, Kamal Chikh Echioukh, Elisabeth Coudert, Beatrice Cuche, Edouard de Castro, Anne Estreicher, Maria L Famiglietti, Marc Feuermann, Elisabeth Gasteiger, Pascale Gaudet, Sebastien Gehant, Vivienne Gerritsen, Arnaud Gos, Nadine Gruaz, Chantal Hulo, Nevila Hyka-Nouspikel, Florence Jungo, Arnaud Kerhornou, Philippe Le Mercier, Damien Lieberherr, Patrick Masson, Anne Morgat, Venkatesh Muthukrishnan, Salvo Paesano, Ivo Pedruzzi, Sandrine Pilbout, Lucille Pourcel, Sylvain Poux, Monica Pozzato, Manuela Pruess, Nicole Redaschi, Catherine Rivoire, Christian J A Sigrist, Karin Sonesson, Shyamala Sundaram, Cathy H Wu, Cecilia N Arighi, Leslie Arminski, Chuming Chen, Yongxing Chen, Hongzhan Huang, Kati Laiho, Peter McGarvey, Darren A Natale, Karen Ross, C R Vinayaka, Qinghua Wang, Yuqi Wang, and Jian Zhang. Uniprot: the universal protein knowledgebase in 2023. *Nucleic Acids Research*, 51:D523–D531, 1 2023.
2. Philippe Le Mercier, Jerven Bolleman, Edouard De Castro, Elisabeth Gasteiger, Parit Bansal, Andrea H. Auchincloss, Emmanuel Boutet, Lionel Breuza, Cristina Casals-Casas, Anne Estreicher, Marc Feuermann, Damien Lieberherr, Catherine Rivoire, Ivo Pedruzzi, Nicole Redaschi, and Alan Bridge. Swissbiopics—an interactive library of cell images for the visualization of subcellular location data. *Database: The Journal of Biological Databases and Curation*, 2022:1–5, 2022.
