## Supplementary File S1 for "Developing *SCL2205*: A Protein Sequence-based Spatial Modelling Dataset for the Protein Language Model Frontier"

### Supplementary Methodology

#### Manual label mapping dictionary

A **Supplementary File S2; Table 1** contains the manual mapping and harmonisation of sub-cellular localisation (SCL) labels.

#### Taxonomic composition

**Table 1. Distribution of data across kingdoms and preparation stages**: *PrpS0* (22152 proteins) and *PrpS1* (19074 proteins; the ***SCL2205*** dataset), captured in **Fig. 2** of the main text. The columns contain the taxonomic classification with data preparation stage sub-columns. The first and second rows are counts and percentages, respectively.

| **Kingdom** | **Metazoa** | | **Viridiplantae** | | **Fungi** | | **Other *** | |
|---|---|---|---|---|---|---|---|---|
| **Prep stage** | *PrpS0* | *PrpS1* | *PrpS0* | *PrpS1* | *PrpS0* | *PrpS1* | *PrpS0* | *PrpS1* |
| **Counts** | 13390 | 10786 | 4250 | 3857 | 4198 | 4137 | 314 | 294 |
| **Percentage** | 60.45% | 56.55% | 19.19% | 20.22% | 18.95% | 21.69% | 1.42% | 1.54% |

* No taxonomy assignment at the *Kingdom* level.

#### Sequence alignments

We used PSI-BLAST for pairwise *all-against-all* alignments and `makeblastdb` to generate search databases, with the parameters outlined below. Unaccounted parameters were used with their default values.

**Database Generation**

```
makeblastdb  -parse_seqids \
    -dbtype ‘‘prot’’ \
    -in <File_fasta> \ # variable  depending on input
    -out <str, in> \ # variable depending on input
    -title <String> # variable depending on input
```

**Pairwise Alignment**

```
psiblast -query <query_file> \ # variable depending on input
    -db <str> \ # variable depending on input
    -out <out_file> \ # variable depending on input
    -evalue 0.005 \
    -word_size <2=<int<=5> \ # variable depending on similarity threshold
    -matrix BLOSUM62 \
    -outfmt ‘‘10 delim=@ qseqid qlen sseqid slen qstart qend sstart send qseq \
            sseq evalue length mismatch gaps pident nident’’ \
    -max_target_seqs 1500 \
    -num_iterations 1 \ # default
    -num_threads <Integer, >=1> # variable depending on the number of cores in the hardware
```

Word size depended on the similarity threshold for homology reduction, similar to the *CD-HIT* tool [1], as follows.

```
if isinstance(p_ident, int):
    if 0 <= p_ident < 50:
        word_size = 2
    elif 50 <= p_ident < 60:
        word_size = 3
    elif 60 <= p_ident < 70:
        word_size = 4
    elif 70 <= p_ident <= 100:
        word_size = 5
```

Throughout, we set the number of iterations to one, effectively using position-specific iterative BLAST (PSI-BLAST) in its BLASTP-equivalent mode. Using more than one iteration would identify distant homologs, which we intentionally avoided. This choice was motivated by two main considerations:

1. The homology paradox – while some relatedness is necessary for learning, excessive similarity can be detrimental. Iterative PSSM can overly reduce homology.

2. Data leakage – including distant homologs during homology augmentation could increase residual overlap boundary between datasets that were initially *overlap-reduced*.

#### Similarity algorithm

We calculated three identity percentages from the resulting alignments using a common numerator – the number of identical amino acids in the alignment – but distinct denominators as follows:

1. Length of the shortest sequence in the pairwise alignment
2. Length of the longest sequence in the pairwise alignment
3. Length of the *no-gap* alignment

Subsequently, for the final percentage similarity score $S$, the weighted geometric mean identity was computed, using the `scipy.stats` module, while penalising the longest sequence by applying its complement weight $(1 - w_l^{seq})$; where $w_l^{seq}$ is the actual weight of the longest sequence. We applied this penalty because percent identity is bounded by the longest sequence, that is, it dictates the maximal possible identity within the alignment *system*. This approach reduces the number of tunable hyperparameters compared to tools such as *CD-HIT*, and eliminates the need for length-based sequence ordering prior to homology reduction.

#### Dataset compositions

##### *Label mapping training-testing preprocessing*

**Table 2.** With and without label mapping independent test set processing before testing. **Original**: Full independent test set size. **Overlap**: Samples remaining after overlap reduction with the training datasets. **Statistics**: Samples remaining after ensuring validity for McNemar's and bootstrapping confidence interval (CI) paired statistical comparison.

| **Sub-cellular location** | **DEEP-SS** | | | **DEEP-HPA** | | |
|---|---|---|---|---|---|---|
| | Original | Overlap | Statistics | Original | Overlap | Statistics |
| **Label Mapping** | | | | | | |
| Secreted† | 522 | 184 | 184 | NA | NA | NA |
| Membrane | 170 | 108 | 108 | 161 | 149 | 149 |
| Mitochondrion | 111 | 74 | 74 | 175 | 154 | 154 |
| Peroxisome | 77 | 30 | 30 | 7 | 7 | 7 |
| Nucleus | 63 | 43 | 43 | 664 | 622 | 622 |
| Cytoplasm | 10 | 6 | 6 | 263 | 249 | 249 |
| ER | 5 | 4 | 4 | 31 | 29 | 29 |
| **Totals** | 1301 | 1210 | 1210 | 1301 | 1210 | 1210 |
| **No Label Mapping** | | | | | | |
| Secreted† | 522 | 189 | 184 | NA | NA | NA |
| Membrane | 170 | 165 | 108 | 161 | 159 | 149 |
| Mitochondrion | 111 | 82 | 74 | 175 | 161 | 154 |
| Peroxisome | 77 | 31 | 30 | 7 | 7 | 7 |
| Nucleus | 63 | 47 | 43 | 664 | 633 | 622 |
| Cytoplasm | 10 | 7 | 6 | 263 | 255 | 249 |
| ER | 5 | 4 | 4 | 31 | 31 | 29 |
| **Totals** | 1301 | 1246 | 1210 | 1301 | 1246 | 1210 |

† Indicates a missing class in the corresponding test split.

*Dataset comparisons training-testing preprocessing*

**Table 3.** *SCL2205*$_s$ and *DEEP-TV*$_s$ versus independent test sets processing, prior to comparison. **Original**: Full independent test set size. **Overlap**: Samples remaining after overlap reduction with the training datasets. **Statistics**: Samples remaining after ensuring validity for McNemar's and bootstrapping CI paired statistical comparison.

| ***SCL2205***$_s$ | | | | | | |
|---|---|---|---|---|---|---|
| **Sub-cellular location** | **DEEP-SS** | | | **DEEP-HPA** | | |
| | Original | Overlap | Statistics | Original | Overlap | Statistics |
| Secreted† | 522 | 184 | 158 | NA | NA | NA |
| Membrane | 170 | 108 | 90 | 161 | 149 | 145 |
| Mitochondrion | 111 | 73 | 64 | 175 | 153 | 149 |
| Peroxisome | 77 | 30 | 24 | 7 | 7 | 7 |
| Nucleus | 63 | 43 | 39 | 664 | 622 | 602 |
| Cytoplasm | 10 | 6 | 6 | 263 | 248 | 238 |
| ER | 5 | 4 | 4 | 31 | 29 | 27 |
| Plastid† | 1 | 0 | 0 | NA | NA | NA |
| **Totals** | 958 | 448 | 385 | 1301 | 1208 | 1168 |
| **SwissProt train-validation (*DEEP-TV*)**$_s$ | | | | | | |
| Secreted† | 522 | 218 | 158 | NA | NA | NA |
| Membrane | 170 | 114 | 90 | 161 | 149 | 145 |
| Mitochondrion | 111 | 77 | 64 | 175 | 163 | 149 |
| Peroxisome | 77 | 37 | 24 | 7 | 7 | 7 |
| Nucleus | 63 | 44 | 39 | 664 | 627 | 602 |
| Cytoplasm | 10 | 7 | 6 | 263 | 244 | 238 |
| ER | 5 | 4 | 4 | 31 | 27 | 27 |
| Plastid† | 1 | 0 | 0 | NA | NA | NA |
| **Totals** | 958 | 501 | 385 | 1301 | 1217 | 1168 |

† Indicates a missing class in the corresponding test split.

*Software and tools information*

**Software specifications**

1. Python: 3.11
2. JupyterLab: 4.1.5
3. BLAST: 2.13.0, build Feb 2 2022 15:38:31
4. RefSeq: BLASTDB Version 5, Oct 26 2024 05:37
5. Pandas: 2.2.1
6. Numpy: 1.26.4
7. Sklearn: 1.4.1.post1
8. Scipy: 1.15.2
9. Matplotlib: 3.10.1
10. Seaborn: 0.12.2
11. Sentencepiece: 0.1.99
12. Torch: 2.8.0+cu128

### Supplementary Results

#### Mapping ablation performance

**Table 4.** Mapping performance by sub-cellular location for DEEP-SS and DEEP-HPA. While we observed a remarkable improvement in label mapping for SwissProt sorting signal (*DEEP-SS*) (9%), the improvement for human protein atlas (*DEEP-HPA*) was marginal.

| Sub-cellular location | Baseline Prevalence | Label Mapping | No Label Mapping |
|---|---|---|---|
| **DEEP-SS** | | | |
| Macro PR-AUC* | 0.143 | 0.296 | 0.206 |
| Secreted | 0.410 | 0.777 | 0.788 |
| Nucleus | 0.096 | 0.555 | 0.242 |
| Membrane | 0.241 | 0.391 | 0.170 |
| Mitochondrion | 0.165 | 0.243 | 0.161 |
| Peroxisome | 0.067 | 0.070 | 0.053 |
| ER | 0.009 | 0.021 | 0.010 |
| Cytoplasm | 0.013 | 0.015 | 0.016 |
| **DEEP-HPA** | | | |
| Macro PR-AUC* | 0.167 | 0.205 | 0.190 |
| Secreted | NA | NA | NA |
| Nucleus | 0.514 | 0.584 | 0.568 |
| Membrane | 0.123 | 0.134 | 0.134 |
| Mitochondrion | 0.127 | 0.206 | 0.160 |
| Peroxisome | 0.006 | 0.011 | 0.010 |
| ER | 0.024 | 0.036 | 0.033 |
| Cytoplasm | 0.206 | 0.260 | 0.233 |

* Overall metric area under the precision–recall curve (PR-AUC). We computed the average prevalence across classes for each test set.

#### Convolutional neural network (CNN) model performance

**Table 5.** *SCL2205$_s$* performance benchmarking against *DEEP-TV$_s$* using two external datasets (*DEEP-SS* and *DEEP-HPA*) based on CNN models. Although *DEEP-TV$_s$* exhibits better discriminative ability (PR-AUC) than *SCL2205* – by 7.7% points on *DEEP-SS* – the difference is marginal for *DEEP-HPA*.

| Sub-cellular location | Baseline Prevalence | *SCL2205$_s$* Eight-class | *DEEP-TV$_s$* Eight-class |
|---|---|---|---|
| **DEEP-SS** | | | |
| **Macro PR-AUC*** | 0.143 | 0.257 | 0.334 |
| Secreted | 0.410 | 0.657 | 0.723 |
| Membrane | 0.234 | 0.384 | 0.353 |
| Mitochondrion | 0.166 | 0.356 | 0.315 |
| Nucleus | 0.101 | 0.291 | 0.737 |
| Peroxisome | 0.062 | 0.074 | 0.096 |
| ER | 0.010 | 0.023 | 0.098 |
| Cytoplasm | 0.016 | 0.014 | 0.014 |
| Plastid | NA | NA | NA |
| **DEEP-HPA** | | | |
| **Macro PR-AUC*** | 0.167 | 0.194 | 0.184 |
| Secreted† | NA | NA | NA |
| Membrane | 0.124 | 0.135 | 0.127 |
| Mitochondrion | 0.128 | 0.147 | 0.147 |
| Nucleus | 0.515 | 0.576 | 0.603 |
| Peroxisome | 0.006 | 0.045 | 0.006 |
| ER | 0.023 | 0.029 | 0.025 |
| Cytoplasm | 0.204 | 0.231 | 0.198 |
| Plastid† | NA | NA | NA |

* Overall metric PR-AUC. We computed the average prevalence across classes for each test set.

† Category missing in dataset.

Protein language model (PLM) model performance

**Table 6.** *SCL2205*$_s$ performance benchmarking against *DEEP-TV*$_s$ using two external datasets (*DEEP-SS* and *DEEP-HPA*) based on PLMs. Although *SCL2205* exhibits better discriminative ability (PR-AUC) than *DEEP-TV*$_s$ – by 10.8% points on *DEEP-SS* – the difference is marginal for *DEEP-HPA*.

| **Sub-cellular location** | **Baseline Prevalence** | ***SCL2205*$_s$ Eight-class** | ***DEEP-TV*$_s$ Eight-class** |
|---|---|---|---|
| | | **DEEP-SS** | |
| **Macro PR-AUC*** | 0.143 | 0.561 | 0.453 |
| Secreted | 0.410 | 0.895 | 0.798 |
| Membrane | 0.234 | 0.876 | 0.751 |
| Mitochondrion | 0.166 | 0.953 | 0.975 |
| Nucleus | 0.101 | 0.959 | 0.557 |
| Peroxisome | 0.062 | 0.197 | 0.060 |
| ER | 0.010 | 0.020 | 0.012 |
| Cytoplasm | 0.016 | 0.023 | 0.021 |
| Plastid† | NA | NA | NA |
| | | **DEEP-HPA** | |
| **Macro PR-AUC*** | 0.167 | 0.345 | 0.359 |
| Secreted† | NA | NA | NA |
| Membrane | 0.124 | 0.430 | 0.489 |
| Mitochondrion | 0.128 | 0.594 | 0.643 |
| Nucleus | 0.515 | 0.818 | 0.761 |
| Peroxisome | 0.006 | 0.012 | 0.030 |
| ER | 0.023 | 0.037 | 0.037 |
| Cytoplasm | 0.204 | 0.180 | 0.193 |
| Plastid† | NA | NA | NA |

* Overall metric PR-AUC. We computed the average prevalence across classes for each test set.
† Category missing in dataset.

#### **SCL2205** final composition summaries

*Training-validation-testing stream composition*

**Table 7.** Sub-cellular location counts across dataset splits.

| Sub-cellular location | Original | Training | Validation | Testing* |
|---|---|---|---|---|
| Membrane | 4718 | 3837 | 277 | 604 |
| Nucleus | 4179 | 3202 | 329 | 648 |
| Secreted | 2635 | 2245 | 131 | 259 |
| Cytoplasm | 2465 | 1922 | 166 | 377 |
| Cytoplasm;Nucleus | 2282 | 1796 | 160 | 326 |
| Mitochondrion | 946 | 740 | 67 | 139 |
| Plastid | 587 | 467 | 37 | 83 |
| Centrosome;Cytoplasm;Cytoskeleton;MTOC** | 308 | 228 | 27 | 53 |
| Cytoplasm;Membrane | 242 | 190 | 17 | 35 |
| Cytoplasm;Cytoskeleton | 215 | 162 | 15 | 38 |
| ER | 185 | 146 | 14 | 25 |
| Cell projection | 176 | 128 | 16 | 32 |
| Peroxisome | 136 | 120 | 4 | 12 |
| **Totals** | **19074** | **15183** | **1260** | **2631** |

* The *Testing* split is common across *cross-validation-testing* (CVT) and *training-validation-testing* (TVT) streams.
** For brevity, Microtubule organizing center (MTOC).

*Cross-validation-testing stream composition*

Table 8: Sub-cellular location counts for Train, Validation, and Test splits across five cross-validation folds.

| Sub-cellular location | Training | Validation | Testing* |
|---|---|---|---|
| **Fold 0** | | | |
| Membrane | 3804 | 310 | 604 |
| Nucleus | 3214 | 317 | 648 |
| Secreted | 2257 | 119 | 259 |
| Cytoplasm | 1916 | 172 | 377 |
| Cytoplasm;Nucleus | 1806 | 150 | 326 |
| Mitochondrion | 744 | 63 | 139 |
| Plastid | 475 | 29 | 83 |
| Centrosome;Cytoplasm;Cytoskeleton;MTOC** | 228 | 27 | 53 |
| Cytoplasm;Membrane | 194 | 13 | 35 |
| Cytoplasm;Cytoskeleton | 161 | 16 | 38 |
| ER | 144 | 16 | 25 |
| Cell projection | 127 | 17 | 32 |
| Peroxisome | 117 | 7 | 12 |
| **Totals** | **15187** | **1256** | **2631** |
| **Fold 1** | | | |
| Membrane | 3836 | 278 | 604 |
| Nucleus | 3217 | 314 | 648 |
| Secreted | 2241 | 135 | 259 |
| Cytoplasm | 1927 | 161 | 377 |
| Cytoplasm;Nucleus | 1784 | 172 | 326 |
| Mitochondrion | 751 | 56 | 139 |
| Plastid | 474 | 30 | 83 |
| Centrosome;Cytoplasm;Cytoskeleton;MTOC** | 223 | 32 | 53 |
| Cytoplasm;Membrane | 191 | 16 | 35 |
| Cytoplasm;Cytoskeleton | 161 | 16 | 38 |
| ER | 153 | 7 | 25 |
| Cell projection | 128 | 16 | 32 |
| Peroxisome | 117 | 7 | 12 |

*Continued on next page*

| Sub-cellular location | Train | Validation | Test |
|---|---|---|---|
| **Totals** | **15203** | **1240** | **2631** |
| **Fold 2** | | | |
| Membrane | 3819 | 295 | 604 |
| Nucleus | 3209 | 322 | 648 |
| Secreted | 2257 | 119 | 259 |
| Cytoplasm | 1918 | 170 | 377 |
| Cytoplasm;Nucleus | 1790 | 166 | 326 |
| Mitochondrion | 740 | 67 | 139 |
| Plastid | 467 | 37 | 83 |
| Centrosome;Cytoplasm;Cytoskeleton;MTOC** | 231 | 24 | 53 |
| Cytoplasm;Membrane | 190 | 17 | 35 |
| Cytoplasm;Cytoskeleton | 165 | 12 | 38 |
| ER | 148 | 12 | 25 |
| Cell projection | 131 | 13 | 32 |
| Peroxisome | 120 | 4 | 12 |
| **Totals** | **15185** | **1258** | **2631** |
| **Fold 3** | | | |
| Membrane | 3817 | 297 | 604 |
| Nucleus | 3227 | 304 | 648 |
| Secreted | 2256 | 120 | 259 |
| Cytoplasm | 1926 | 162 | 377 |
| Cytoplasm;Nucleus | 1796 | 160 | 326 |
| Mitochondrion | 741 | 66 | 139 |
| Plastid | 469 | 35 | 83 |
| Centrosome;Cytoplasm;Cytoskeleton;MTOC** | 228 | 27 | 53 |
| Cytoplasm;Membrane | 190 | 17 | 35 |
| Cytoplasm;Cytoskeleton | 161 | 16 | 38 |
| ER | 149 | 11 | 25 |
| Cell projection | 131 | 13 | 32 |
| Peroxisome | 119 | 5 | 12 |
| **Totals** | **15210** | **1233** | **2631** |
| **Fold 4** | | | |
| Membrane | 3861 | 253 | 604 |
| Nucleus | 3221 | 310 | 648 |
| Secreted | 2280 | 96 | 259 |
| Cytoplasm | 1922 | 166 | 377 |
| Cytoplasm;Nucleus | 1814 | 142 | 326 |
| Mitochondrion | 739 | 68 | 139 |
| Plastid | 461 | 43 | 83 |
| Centrosome;Cytoplasm;Cytoskeleton;MTOC** | 227 | 28 | 53 |
| Cytoplasm;Membrane | 193 | 14 | 35 |
| Cytoplasm;Cytoskeleton | 161 | 16 | 38 |
| ER | 143 | 17 | 25 |
| Cell projection | 127 | 17 | 32 |
| Peroxisome | 116 | 8 | 12 |
| **Totals** | **15265** | **1178** | **2631** |

* The *Testing* split is common across CVT and TVT streams.
** For brevity, Microtubule organizing center (MTOC).
